# Mast cells initiate lymphocyte egress from distant lymph nodes upon skin inflammation via a RANKL–sphingosine-1-phosphate axis

**DOI:** 10.64898/2026.06.24.734311

**Authors:** Konstantinos Katsoulis-Dimitriou, Waqar Umer, Ali El-Bizri, Laura Knop, Tanja Schickschneit, Aaron Hoffman, Lea M. Schmitter, Kathleen Baumgart, Nouria Jantz-Naeem, Vladyslava Dovhan, Charlotte Heidelbach, Lars Philipsen, Andreas J Müller, Sascha Kahlfuss, Thomas Schüler, Stephan Fricke, Jan Dudeck, Anne Dudeck

## Abstract

Receptor activator of NFκB ligand (RANKL) is important for bone metabolism, but also modulates immune processes. We showed that mast cells (MCs) are involved in RANKL regulation, but the importance of MC-derived RANKL in skin inflammation has not yet been investigated. In contact hypersensitivity (CHS), the absence of MC-derived RANKL led to reduced skin inflammation due to impaired leukocyte infiltration and blood lymphopenia. Surprisingly, we observed a massive hyperplasia of the distant inguinal lymph nodes in the absence of MC-RANKL. Using adoptive transfers, flow cytometry and whole-mount 3D imaging, we demonstrated that this was not caused by structural maladaptation, but rather by the inability of lymphocytes to exit in a timely manner. Importantly, RANKL deletion in skin MCs only replicated the effect of LN hyperplasia and blood lymphopenia. Moreover, MCs were involved in serum sphingosine-1-phosphate (S1P) regulation during sensitization and challenge. Intravascular administration of S1P restored timely lymphocyte egress, demonstrating a MC-induced organ-spanning RANKL-S1P axis. Consequently, peripheral skin MC-derived RANKL is essential for the timely lymphocyte egress from distant LNs, which may have important implications for the targeted treatment of inflammatory skin diseases.

## Introduction

Receptor activator of NFκB ligand (RANKL) and its receptor RANK are members of the TNF super family and best known for their role in skeletal development and bone homeostasis. RANKL stimulates osteoclastogenesis and thus bone resorption, and thereby also promotes bone loss during age-related osteoporosis and inflammatory diseases (1, 2). However, there is increasing evidence for immune regulating RANKL functions. As reviewed by Mueller and Hess, RANKL is involved in lymph node (LN) formation, since mice lacking RANKL or RANK expression completely lack LNs and show abnormalities in secondary lymphoid organs (SLOs). In detail, RANKL released by lymphoid tissue organizer cells (LTO) is responsible for recruitment of lymphoid tissue inducer cells (LTI), which is crucial for LN formation (3). Moreover, RANKL initiates SLO development, and promotes growth of LNs, peyer’s patches and other SLO (4, 5). Of note, RANKL has also been shown to modulate skin inflammation. Upon UV irradiation, RANKL released by keratinocytes interacts with RANK on Langerhans cells and dermal dendritic cells (DCs) and promotes T cell activation and Treg expansion (6, 7).

Mast cells (MCs) are long-lived, tissue-resident innate immune cells populating predominantly interface tissues, such as the skin. Due to their immediate response to IgE/FcεRI-crosslinking, MCs are best known as key effector cells in type I allergic reactions. MCs contain a huge amount of secretory granules that are filled with preformed mediators and instantaneously released by degranulation, which is later followed by release of *de novo* synthesized mediators (8). However, MCs also sense pathogen patterns, inflammatory stimuli and cell stress signals, such as IL-33 and extracellular ATP (9, 10) and play important roles in infections and (non-allergic) inflammatory diseases as well (11). Moreover, MCs drive innate immune response and modulate adaptive immunity by direct communication with other immune cells (12). For example, we demonstrated a bidirectional interaction between MCs and DCs, where DCs transfer MHCII complexes to MCs (13), while in turn engulfing intact MC granules, which promotes DC maturation, migration to draining LN (dLN) and their T cell priming efficiency (14). Recently, we proved that perivascular skin MCs form intravascular sheets, and degranulate directionally in the bloodstream. Preformed TNF, embedded in MC granules and infused via this route into the bloodstream was crucial for priming blood circulating neutrophils to license their migration to the inflamed skin (15).

We have shown that MCs are involved in the increase of serum RANKL levels upon bone fracture healing and promote osteoclast formation (16). Using an MC-specific conditional knockout, we have also demonstrated that MC-derived RANKL is involved in ovariectomy-induced osteoporosis (17). Considering the high density and sentinel function of MCs in the skin, we here examined the role of MC-derived RANKL in skin inflammation. We found that the release of RANKL by skin MCs is crucial for skin inflammation, but surprisingly, by initiating lymphocyte egress from distant inguinal LNs, which is mandatory for their migration into the skin and pro-inflammatory effects.

## Results

We generated a conditional RANKL knockout mouse line by crossing RANKL^fl/fl^ and Mcpt5-Cre mice (17, 18), where Cre expression is specific for connective tissue type MC. Efficient abrogation of RANKL expression has been confirmed *in vivo* in ear skin MCs and *in vitro* in peritoneal cultured MCs of Mcpt5-Cre^+^ RANKL^fl/fl^ (MC^ΔRANKL^) as compared to Cre^-^ control mice (MC^wt^) **(Suppl. Figure 1A)** (17). Importantly, both MC numbers and distance to blood vessels were not altered upon RANKL deletion **(Suppl. Figure 1B).** MC ability to degranulate upon stimulation with calcium Ionophore, ATP, ATP and IL-33, or crosslinking of the IgE receptor was also unaltered **(Suppl. Figure 1C).** Given the relevance of RANKL for lymphoid tissue development, we analyzed LN and blood immune cell populations in steady state. Immunophenotyping showed no difference between MC^ΔRANKL^ and MC^wt^ in LN and blood total leukocyte counts and numbers of CD4^+^ and CD8^+^ T cells, and B cells in the steady state **(Suppl. Figure 2A/B)**. Lymphatic tissue that has normally higher activity in the baseline, such as mesenteric LNs and peyer’s patches, showed no differences in T cell memory, effector and naïve populations as well as in total and marginal zone B cells **(Suppl. Figure 2C/D)**.

### Impaired skin inflammation in absence of MC-derived RANKL

The role of MC-derived RANKL in skin inflammation was investigated using the contact hypersensitivity (CHS) mouse model, which is a T cell-mediated delayed type hypersensitivity response. Sensitization with the hapten DNFB on back skin results in hapten-specific T cell expansion in inguinal lymph nodes (LN_in_), which then infiltrate the ear skin upon DNFB challenge and initiate skin inflammation. Importantly, both the ear swelling and the infiltration of leukocytes was massively impaired in MC^ΔRANKL^ as compared to MC^wt^ mice 24h post challenge, particularly of CD8^+^ T cells and neutrophils **(Figure 1A/B)**. At the same time, numbers of all blood lymphocyte populations, CD4^+^ and CD8^+^ T cells, and B cells, were significantly reduced to half in MC^ΔRANKL^ mice as compared to the MC^wt^ control, while neutrophils numbers were not **(Figure 1C/D)**. Upon first hapten encounter on back skin, skin DCs process haptenized self-proteins and migrate to draining LN_in_ to prime hapten specific T cells (12). Given our previous observations on MC-DC communication (14, 19), we questioned whether the reduced skin and blood lymphocyte numbers may result from impaired DC migration. Using FITC-induced CHS allowing the tracking of FITC^+^ DC (Dudeck et al 2011), we found that both the numbers of total DCs and of FITC^+^ DCs in LN_in_ were not altered in MC^ΔRANKL^ as compared to MC^wt^ mice **(Figure 1E)**.

**Figure 1:**
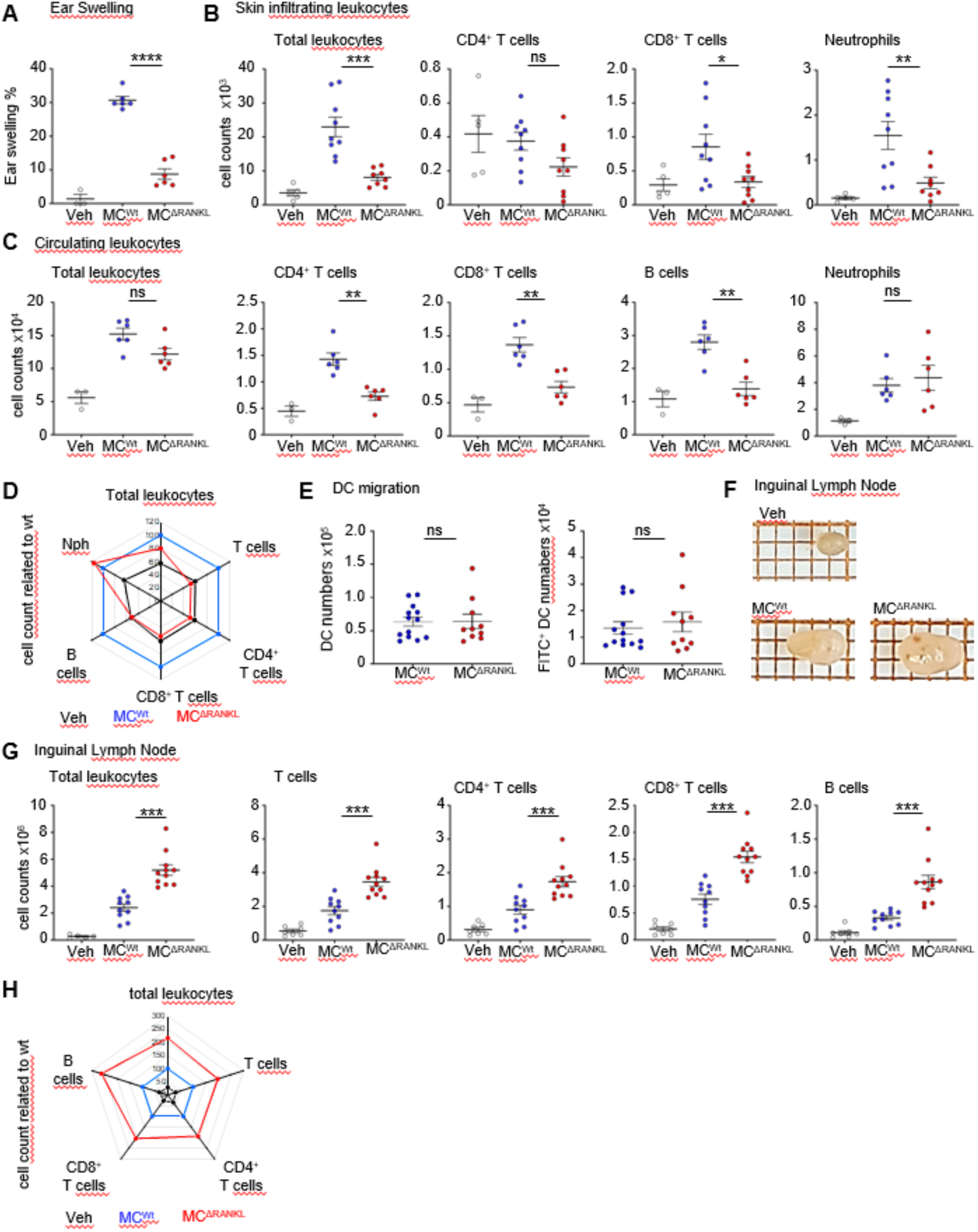
Absence of MC-derived RANKL results in blood lymphopenia but inguinal LN hyperplasia upon CHS elicitation. Mice were sensitized on the back skin and after 6 days challenged on the ear skin for 24h. (**A**) Ear skin swelling response. (**B**) Total numbers of lymphocytes infiltrating the ear skin. (**C**) Blood lymphocyte numbers per 100 μl of blood. (**D**) Radar plot showing blood lymphocyte numbers per 100 μl of blood, MC^wt^ is set as the baseline. (**E**) MC^ΔRANKL^ or MC^wt^ mice were sensitized by epicutaneous application of 100 µl fluorescein isothiocyanate (FITC) on the shaved back. 48h after sensitization, LN_in_ were isolated and total numbers of DCs (left) and FITC^+^ DCs, that have migrated from the skin (right) were analyzed by flow cytometry. (**F**) Side by side comparison of whole LN_in_ from MC^ΔRANKL^ and MC^wt^ and veh mice. (**G**) Lymphocyte numbers of the inguinal LN. (**H**) Radar plot showing total numbers of the inguinal LN, normalized by setting MC^wt^ as 100%. All graphs are depicted as mean and SD. **** p < 0.0001; *** p < 0.001; ** p < 0.01; * p < 0.05.

### Lack of MC-derived RANKL leads to LN_in_ hyperplasia upon CHS elicitation

Surprisingly, we found that LN_in_ draining the sensitized back skin exhibited a dramatic hyperplasia at 24h after challenge **(Figure 1F)**. The total numbers of CD3^+^ T cells, CD4^+^ T cells, CD8^+^ T cells, and B cells were massively increased in MC^ΔRANKL^ as compared to MC^wt^ mice **(Figure 1G).** By normalizing the number of lymphocytes in MC^ΔRANKL^ mice to MC^wt^ mice we found that all lymphocyte populations were at least doubled **(Figure 1H)**.

Given the importance of RANKL for LN anlagen, development and growth, we aimed to evaluate any structural dysregulation or maladaptation of LN_in_ in context to the LN hyperplasia. Therefore, we established a whole mount staining of the entire intact LN_in_, modified according to Li et al. (20), followed by tissue clearing and two-photon microscopy of the entire cleared LNs **(suppl. Figure 3A)**. First, LN 3D projection and optimal sectioning in the midpoint of 3D images proved respective whole-mount staining efficiency and specificity of T and B cell areas, blood vessels and lymph vessels **(suppl. Figure 3B/C)**. In order to not only visualize the respective zones but also quantify structural parameters, the whole LNs were scanned and LN volume as well as T cell and B cell areas were reconstructed using the Imaris surface tool with machine learning function, based on fluorescent intensity and quantified as volumes **(Figure 2B-D)**. The size of the whole LN_in_, as well as the T cell and B cell zones separately, was found to be consistently increased, confirming the significant LN hyperplasia and increase of all lymphocyte subsets in absence of MC-derived RANKL **(Figure 2B-D)**. However, the ratios of T cell to B cell zone volume, as well as of T cell and B cell zones to the total LN_in_ volume, were not significantly different, showing that the LN_in_ was generally enlarged without a specific impact to some compartments **(Figure 2E).** Using the filament tracer for blood vessel reconstruction and quantification, we found that vessel volume and diameter was not increased in MC^ΔRANKL^ while total segment length was increased, showing more outstretched vessels correlating to the increased LN size **(Figure 2F)**. Overall, no visible structural defects or maladaptation were observed except for the LN enlargement.

**Figure 2:**
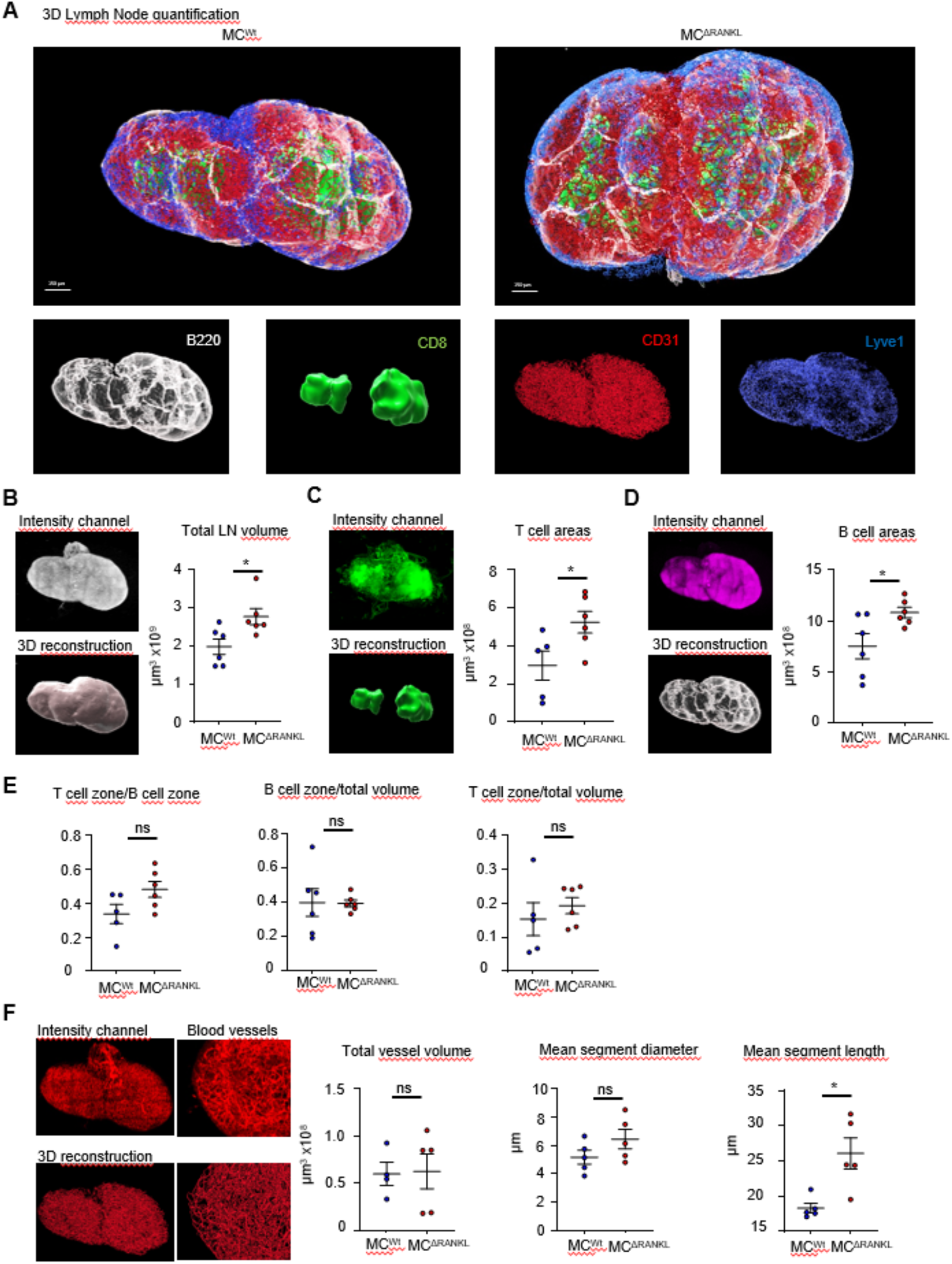
LN hyperplasia in MC^ΔRANKL^ mice is not driven by structural maladaptation. (**A**) Whole-mount staining, 3D imaging and reconstruction of LN_in_ from MC^wt^ and MC^ΔRANKL^ mice, at 24h after hapten challenge, including B cell (B220) and T cell zones (CD8), blood vessels (CD31) and lymphatic vessels (Lyve1). B. Total LN_in_ volume (**B**) was measured by creating a surface based on all channels with Imaris machine learning, and is plotted in the cumulative graph. C/D. T cell (**C**) and B cell (**D**) area volume was measured by creating a surface with Imaris and plotted in the cumulative graph. (**E**) B cell and T cell areas are shown as ratio with each other and with the total volume of the LN_in_. (**F**) Blood vessels were reconstructed with the Imaris filament tracer. Cumulative graphs of blood vessel volume, segment diameter and length. All graphs are depicted as mean and SD.

Given how important fibroblastic reticular cells (FRCs) are for the development and expansion of the LN we investigated whether steady state alterations existed upon MC-RANKL deletion (21). LN_in_ were tested for FRC related chemokine expression by qPCR. The main FRC homeostatic chemokines *Ccl19* and *Ccl21*, as well as *Cxcl9*, *Cxcl10* and *Cxcl13* were unaltered in MC^ΔRANKL^ mice **(suppl. Figure 4A)**. To address whether LN enlargement reflected a broader pre-existing structural maladaptation, we performed additional multi epitope ligand cartography (MELC) analysis of lymph-node organization. This included the spatial localization of major lymphocyte populations in relation to the FRC network, with quantitative assessment of the distances of CD4⁺ T cells, CD8⁺ T cells, and B cells from FRCs, as well as their relative cellular frequencies within the LN. Importantly, none of these parameters differed between groups. Thus, despite the clear increase in LN size, the relative distribution of major lymphocyte populations, their positioning with respect to the FRC scaffold, and their overall representation within the tissue remained preserved. Together with the unchanged expression of key lymphoid chemokines, these data argue against a pre-existing defect in LN stromal organization or lymphocyte compartmentalization as the cause of the observed LN hyperplasia. **(suppl. Figure 4B/C)**.

To investigate whether the LN hyperplasia is driven by a lymphocyte intrinsic disability to expand or to migrate, lymphocytes were isolated from spleen and LNs from sensitized wt mice at day 6 post DNFB, CFSE labeled and adoptively transferred into sensitized MC^ΔRANKL^ and MC^wt^ mice at 1 day after DNFB sensitization of recipient mice. 5 days later, recipient mice LN_in_ were analysed **(Figure 3A)**. As expected, endogenous (CFSE^-^) lymphocyte numbers were markedly increased in MC^ΔRANKL^ compared to MC^wt^ mice, excluding any transfer-induced effects on LN_in_ hyperplasia **(Figure 3B).** Of note, we observed a massively increased number of transferred CFSE^+^ wt lymphocytes in LN_in_ of MC^ΔRANKL^ recipient mice compared to MC^wt^ mice, again affecting all lymphocyte populations, CD4^+^ and CD8^+^ T cells and B cells, indicating that the transferred wt lymphocytes reach the LN_in_ and undergo expansion, but are retained in absence of MC-derived RANKL **(Figure 3C).** Collectively, these data demonstrates that the LN hyperplasia is not driven by lymphocyte intrinsic effects but dictated by the tissue without any developmental or structural LN defects.

**Figure 3:**
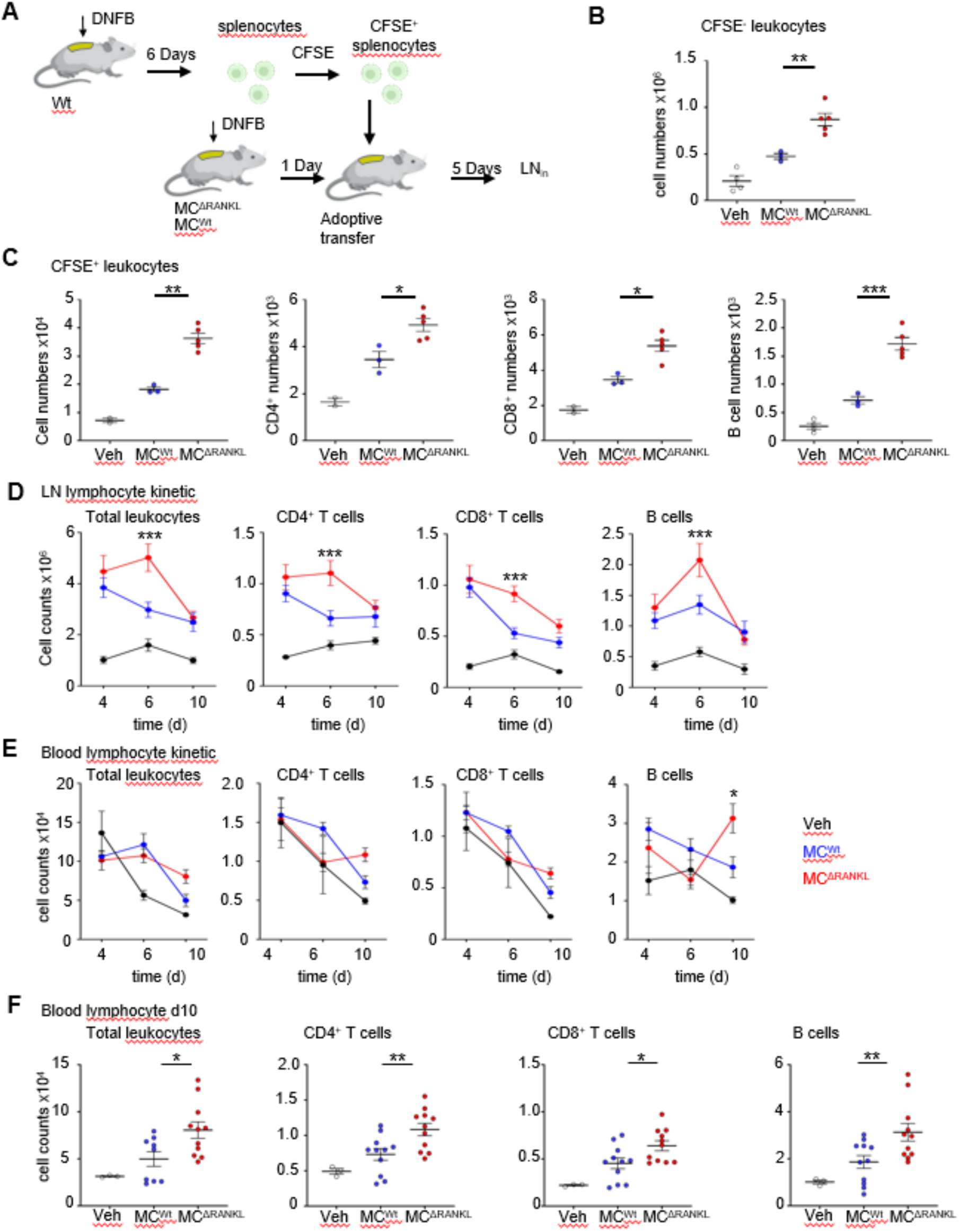
LN_in_ hyperplasia in MC^ΔRANKL^ mice is transient and driven by tissue effects. (**A**) MC^ΔRANKL^ or MC^wt^ mice, as well as wt mice were sensitized on the back skin for 6 days. Lymphocytes were isolated from spleens and LNs of sensitized wt mice, labeled with CFSE and adoptively transferred into MC^ΔRANKL^ or MC^wt^ mice 1 day after sensitization. 5 days later, flow cytometric analysis of LN lymphocytes was performed. (**B**) Total numbers of lymphocytes in the inguinal lymph node. (**C**) Numbers of CFSE^+^ labeled lymphocyte subsets in the inguinal lymph node of recipient mice. (**D**) Mice were sensitized on the back skin with DNFB and total leukocyte numbers were evaluated by flow cytometry on days 4, 6 and 10. Kinetic assay of inguinal LN lymphocyte numbers at days 4, 6 and 10 after sensitization. (**E**) Kinetic assay of blood lymphocyte numbers, per 100μl of blood, at days 4, 6 and 10 after sensitization (**F**) Blood lymphocyte numbers per 100μl of blood at day 10 after sensitization. All graphs are depicted as mean and SD. group **** p < 0.0001; *** p < 0.001; ** p < 0.01; * p < 0.05.

### Transient LN hyperplasia and blood lymphopenia persistently impair skin inflammation

Next, the kinetics of blood lymphopenia and LN_in_ hyperplasia were determined at days 4, 6, and 10 after sensitization and were found to be transient (**Figure 3D/E).** More specifically, LN_in_ showed increased cellularity at day 4, with no differences yet between MC^ΔRANKL^ and MC^wt^ mice. On day 6, the highest lymphocyte count was reached, with the hyperplasia in MC^ΔRANKL^ mice of almost twofold more lymphocytes than in MC^wt^ mice. On day 10, LN_in_ cellularity was reduced to the same level in both MC^wt^ mice and MC^ΔRANKL^ mice compared to day 4, indicating lymphocyte egress **(Figure 3D).** Transient effects were observed in the blood as well, since lymphocyte numbers were similar between MC^ΔRANKL^ and MC^wt^ mice at day 4 but reduced at day 6 **(Figure 3E).** Moreover, the blood lymphopenia was reversed by day 10, with significantly more CD4^+^ T and CD8^+^ T cells, and B cells in the blood of MC^ΔRANKL^ mice compared to MC^wt^ mice **(Figure 3F)**, indicating a delayed onset of lymphocyte egress in the absence of MC-derived RANKL. Given the delayed lymphocyte egress in MC^ΔRANKL^ mice, also kinetics of skin inflammation and ear skin lymphocyte infiltration after challenge were examined **(Figure 4A/B)**. In MC^wt^ mice, the ear swelling peaks at approximately 40% at 48h after challenge with a rapid decline thereafter. In MC^ΔRANKL^ mice, the ear swelling was significantly reduced in the first 48h compared to MC^wt^ mice, with a peak of only ∼20% at day 3 and declined thereafter similarly to MC^wt^ mice **(Figure 4A).** Surprisingly, the reduced lymphocyte infiltration at 24h (see Figure 1A), was reversed at 48h, thereby showing significantly higher numbers of CD4^+^ and CD8^+^ T cells in ear skin **(Figure 4C).** This finding is in line with the reversed lymphocyte levels in the blood at later time points, but contrasting to the impaired ear swelling. We therefore specifically determined the extravasation of lymphocytes to exclude artifacts of blood microvessel lymphocytes instead of those in skin tissue. Blood circulating cells were first stained *in vivo* by i.v injection of PE-conjugated anti-CD45 mAb, while following ear skin digestion, all leukocytes were counterstained with PercP-Cy5.5-conjugated anti-CD45 mAb **(Figure 4C)**. Hence, skin lymphocyte numbers were increased in MC^ΔRANKL^ compared to MC^wt^ mice **(Figure 4D)**, the extravasation capacity determined as ratio of circulating and tissue lymphocytes was not altered **(Figure 4E)**. In contrast to increased skin lymphocyte numbers, the ear skin concentration of the cytokines IFNγ, TNF, IL-4 and IL-2 were not altered in MC^ΔRANKL^ mice compared to MC^wt^ **(Figure 4F)**.

**Figure 4:**
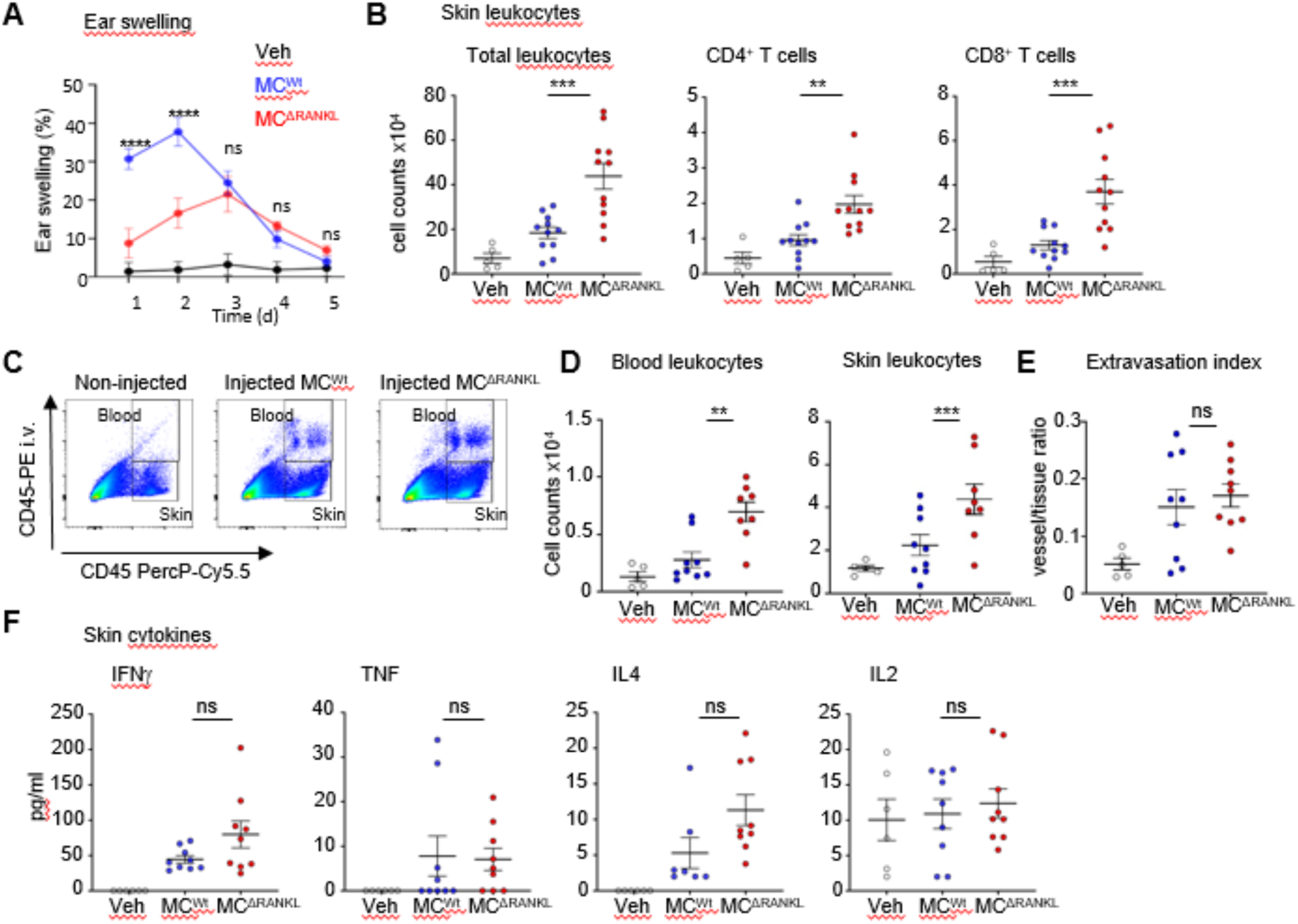
Delayed onset of leukocyte skin infiltration leads to reduced inflammatory symptoms. Mice were sensitized on the back skin and challenged 6 days later on the ear skin for 48h. (**A**) Kinetics of the ear swelling response up to day 5 after DNFB challenge. (**B**) Total numbers of lymphocytes infiltrating the ear skin 48h after challenge. (**C**) Example FACS plot showing the gating strategy for separating vessel lymphocytes to tissue lymphocytes. MC^ΔRANKL^ or MC^wt^ were sensitized and challenged 6 days later on the ear skin. CD45-PE antibody was injected intravenously 48h after challenge, in order to stain all circulating leukocytes. After ear skin digestion, circulating PE-labeled leukocytes were quantified in relation to total leukocytes, stained ex vivo by CD45-PerCP-Cy5.5, to separate lymphocytes in the tissue from the cells in the vessels. (**D**) Graphs showing lymphocyte numbers in the blood vessels and ear skin of the ears. (**E**) Cumulative graph showing extravasation as a normalized fraction of cells in the vessels to cells in the tissue. (**F**) Inflammatory cytokine concentrations in the ear skin were measured 48h after challenge. All graphs are depicted as mean and SD. **** p < 0.0001; *** p < 0.001; ** p < 0.01; * p < 0.05.

### Peripheral skin MC production of RANKL is crucial for lymphocyte egress from distant LNs

Given the LN hyperplasia in absence of MC-RANKL and the strong expression of RANKL by MCs in the steady state **(Figure 5A)**, we investigated whether MCs in skin or LNs are the key source. MCs were depleted only locally in the ear skin of Mcpt5-Cre iDTR mice by i.d. diphtheria toxin (DT) injection (Dudeck et al. 2011), and reconstituted with MCs from either MC^ΔRANKL^ or MC^wt^ mice. Efficient MC depletion and reconstitution was proven by microscopy (data not shown), and CHS was performed 2 weeks after reconstitution. Consequently, only MCs in the DNFB-challenged ear skin lacked RANKL production, while all other MCs, in sensitized back skin and LNs as well, are RANKL-proficient **(Figure 5B)**. Intriguingly, absence of RANKL in ear skin MCs only resulted in reduced ear swelling upon DNFB challenge **(Figure 5C)**, accompanied by impaired infiltration of CD4^+^ and CD8^+^ T cells, and neutrophils **(Figure 5D)**. Moreover, mice lacking RANKL in only ear skin MCs still exhibited the massive LN_in_ hyperplasia with increased numbers of all CD4^+^ and CD8^+^ T cells, and B cells at 24h after challenge, and simultaneous blood lymphopenia with significant reduction in CD4^+^ and CD8^+^ T cell numbers, in comparison to mice reconstituted with wt PCMCs, as observed in MC^ΔRANKL^ **(Figure 5E/F)**. This data indicates that RANKL release by ear skin MCs upon CHS challenge drives a long-range effect towards the distant back skin-draining LN_in_ that is a prerequisite for timely lymphocyte egress.

**Figure 5:**
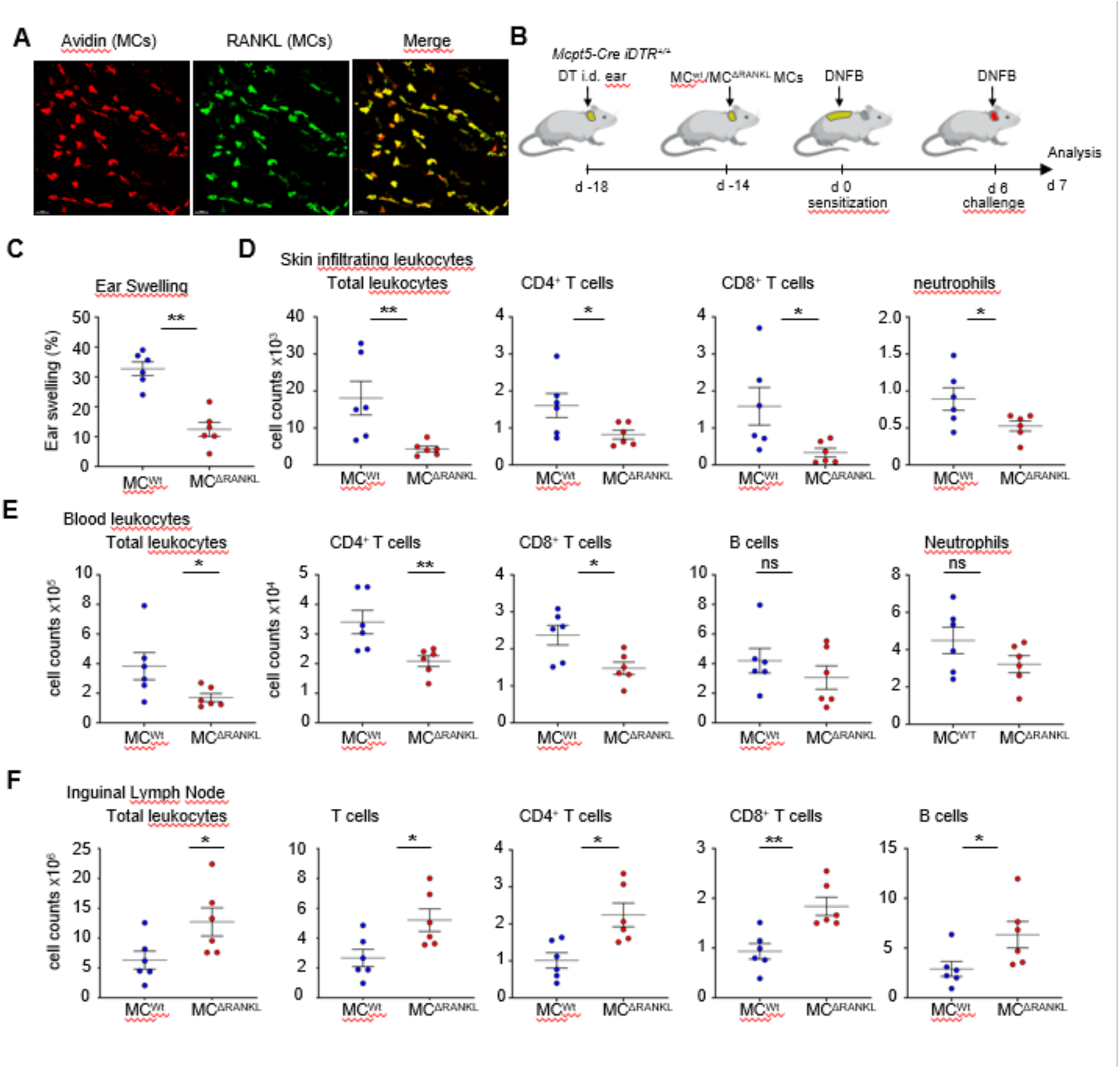
RANKL release by peripheral skin MCs is crucial for lymphocyte egress from distant LN_in_. (**A**) Confocal microscopy of whole mount dermal sheets showing RANKL expression by skin MCs in the steady state (**B**) MCs were depleted only locally in the ear skin by i.d. DT injection in Mcpt5-Cre iDTR mice. Subsequently, ear skin MCs were reconstituted with peritoneally cultured MCs either from MC^ΔRANKL^ or MC^wt^ mice, followed by sensitization and challenge with DNFB. After 24h lymphocyte numbers were validated with flow cytometry. (**C**) Ear skin swelling response. (**D**) Total numbers of lymphocytes infiltrating the ear skin 24h after challenge. (**E**) Blood lymphocyte numbers per 100μl of blood as quantified by flow cytometry. (**F**) Total numbers of lymphocytes in LN_in_ at 24h after challenge. All graphs are depicted as mean and SD. **** p < 0.0001; *** p < 0.001; ** p < 0.01; * p < 0.05.

### MC-derived RANKL triggers lymphocyte egress by inducing an early blood Sphingosine-1-phosphate (S1P) burst

Next, we aimed at deciphering the mechanism underlying the remote RANKL effect from ear skin to distant LN_in_. RANKL has been reported to stimulate the secretion of Sphingosine-1-phosphate (S1P) (22), a signaling molecule initiating lymphocyte migration (23). Since lymphatic endothelial cells (LECs) are known to be an important source of S1P, we next investigated whether RANKL can stimulate S1P release by LECs. To address this, cultured ear skin LECs were treated either with recombinant RANKL or with supernatants from activated PCMCs. PCMC supernatants were collected after either early activation for 2 h or prolonged activation for 24 h. Both recombinant RANKL and supernatants from activated MC^wt^ PCMCs induced S1P release by LECs. In contrast, supernatants from activated MC^ΔRANKL^ PCMCs failed to induce this response. These findings suggest that MC-RANKL is sufficient, and required within the mast cell supernatant, to promote S1P release by LECs. **(Figure 6A).** Moreover, although similar in the steady state, serum S1P steadily increased up to day 4 after sensitization in MC^wt^ mice and to a significantly lesser degree in MC^ΔRANKL^ mice **(Figure 6B)**. This indicates that MC RANKL is involved in the regulation of the S1P response during CHS. By analyzing the kinetics of serum S1P levels upon CHS challenge, we found an early onset of S1P release only 2h after DNFB challenge with a peak at 24h, and decline thereafter until 48h. Importantly, the early S1P production 2h after DNFB compared to veh treated mice was completely missing in MC^ΔRANKL^ mice, with levels even below the veh control group (unspecific skin irritation), while the delayed onset was reduced faster towards significant lower levels at 48h **(Figure 6B).** Surprisingly, intradermal injection of RANKL alone was not sufficient to induce the S1P burst at 2h, indicating a preconditioning during sensitization, a synergistic effect of MC-RANKL with other mediators, or a stabilization of RANKL within MC secretory granules and focused degranulation being involved **(Figure 6C)**. Nevertheless, since the early S1P burst at 2h post DNFB in MC^wt^ mice may be triggered by the immediate release of preformed MC-derived RANKL, we speculated that the phenotype of LN_in_ hyperplasia and blood lymphopenia in MC^ΔRANKL^ mice could be rescued by i.v. injection of recombinant S1P at 2h post DNFB **(Figure 6D)**. Hence, CHS was performed in MC^wt^ and MC^ΔRANKL^ mice, and MC^ΔRANKL^ received an i.v. injection of 500ng recombinant S1P (simulating the S1P concentration in MC^wt^ mice) or PBS as a control at 2h post DNFB, and were compared to MC^wt^ mice. Intriguingly, the lymphocyte infiltration into the ear skin was increased significantly as compared to PBS control mice, reaching the MC^wt^ mice levels **(Figure 6E)**. In line, the single S1P injection was sufficient to significantly reverse both the LN_in_ hyperplasia and blood lymphopenia of CD4^+^ and CD8^+^ T cells, and B cells at 24h after challenge to the MC^wt^ levels **(Figure 6E/F)**. These results reveal that RANKL released peripherally by skin MCs is inducing lymphocyte egress in distant LNs upon skin inflammation through a remote mechanism involving early S1P release into the blood stream.

**Figure 6:**
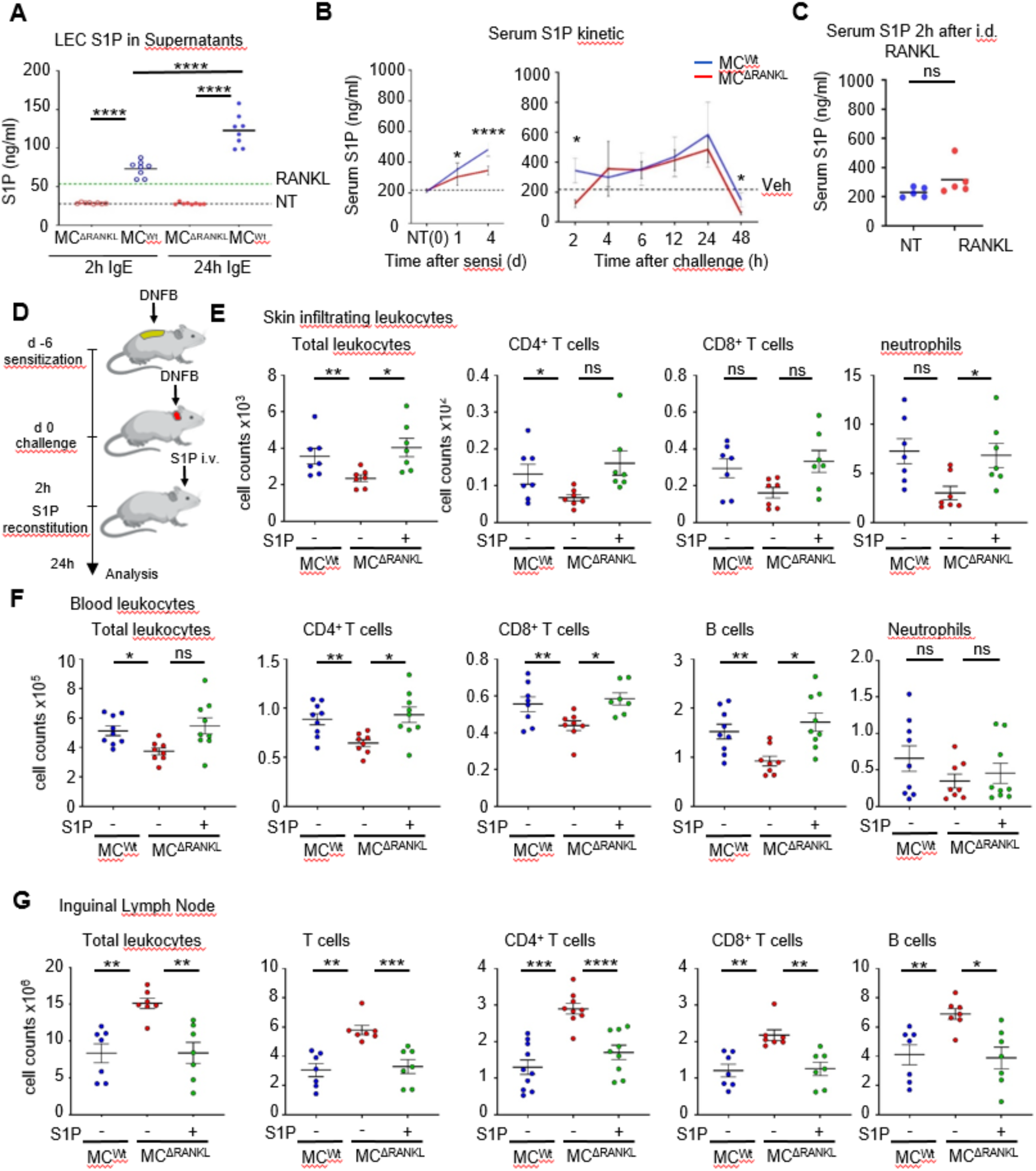
Sphingosine-1-phosphate (S1P) administration rescues LN lymphocyte retention in MC^ΔRANKL^ mice. (**A**) Lymphatic endothelial cells (LECs) were sorted from ear skin and expanded in vitro, then treated with PCMC supernatants that were either activated for 2h or 24h via the FceR in vitro (anti DNP-IgE 24h followed by DNP BSA). LECs were also treated with recombinant RANKL (green line) or were untreated (black line). S1P expression was measured in the supernatants. (**B**) MC^ΔRANKL^ or MC^wt^ were DNFB sensitized and challenged after 6 days, and a kinetic assay of Sphingosine-1-phosphate (S1P) was performed up until 48h after challenge. (**C**) Recombinant RANKL was injected i.d. in the ear and 2h after serum S1P was measured. (**D**) 2h after challenge 500ng of S1P (+) or PBS only as control (-) was injected i.v. Flow cytometric measurements of skin, blood and LN lymphocytes were performed 24h after challenge. (**E**) Total numbers of lymphocytes infiltrating the ear skin. (**F**) Blood lymphocyte numbers per 100μl of blood. (**G**) Total numbers of lymphocytes in LN_in_. All graphs are depicted as mean and SD **** p < 0.0001; *** p < 0.001; ** p < 0.01; * p < 0.05.

## Discussion

There is increasing evidence that peripheral MCs are involved in LN hyperplasia in inflammatory conditions (12). For example, TNF released from skin MCs upon bacterial challenge is rapidly detectable in draining LNs (25, 26). Moreover, MCs play a pivotal role in TNF-independent LN expansion upon intradermal bacterial peptidoglycan injection (26, 27). Kunder et al. showed that after peripheral MC degranulation, released MC granules can enter the lymphatics and reach the draining LN thereby eliciting profound LN hypertrophy, while no MC activation within the LN was observed (28). In addition, we could show that, upon LPS- or hapten-induced skin inflammation, released intact MC granules are taken up by skin resident DCs, which promotes DC maturation, migration to draining LNs and thereby T cell priming (29). Hence, mechanistically, MC-DC communication within the skin, involving both MC granule effects and physical intercellular interaction (13), as well MC mediator drainage from skin through lymphatics contribute to remote LN hyperplasia. However, MCs release a plethora of proinflammatory mediators which, in addition to TNF, may have an impact on LN conditioning in one way or another. We and others have proven that MCs are a relevant source of RANKL (30, 31). However, although RANKL overproduction in hair follicles has been found to cause significant LN hyperplasia (32), the role of MC-derived RANKL in adaptive skin inflammation remains to be investigated.

In this study, we show for the first time that skin MC-derived RANKL is involved in the conditioning of distant LNs upon skin inflammation. While the local inflammatory response is impaired upon hapten challenge in absence of MC-derived RANKL, inguinal LNs show a massive hyperplasia, involving all the lymphocyte populations that expand upon hapten sensitization. Notably, this effect was observed after hapten challenge on ear skin but in the distant inguinal LN_in_, draining the sensitized back skin. Importantly, we could prove that peripheral skin MCs, but not MCs in the LNs, are responsible for this long-distance communication phenomenon by a local ear skin MC depletion and reconstitution with either wt or RANKL deficient MCs. A long distant effect was further supported by the finding that DC migration to LN_in_ upon back skin sensitization was not affected. The massive LN_in_ hyperplasia was accompanied by simultaneous blood lymphopenia, indicating that lymphocytes expanded as expected but fail to egress from LN_in_ to the blood. This could be caused by direct MC-RANKL effects on the LN or by indirect effects via cellular mediators.

Skin MCs are well known to initiate recruitment of innate, but also adaptive immune cells such as T cells (8). For example, direct MC degranulation into the bloodstream is crucial to prime and recruit neutrophils in CHS (15). Hence, MC granules may reach distant organs via the bloodstream and therefore exert long range effects. The plethora of MC mediators, in conjunction with their ability to promote adaptive immune cell expansion by their effects on DC migration and maturation make it difficult to discriminate between their effects on T cell priming vs. recruitment. To exclude MC effects on T cell priming or T cell intrinsic inabilities to egress from LNs, we adoptively transferred wt T cells to mice lacking MC-RANKL and found them to reach the LN_in_, expand properly, but being unable to leave the LN_in_ as well. This finding supported the hypothesis of a missing egress signal, originating from peripheral MCs, that is independent of the T cell priming and expansion capacity.

Studying the kinetics of the inflammatory response, we noticed a lymphocyte egress from distant LN_in_ at later timepoints coinciding with a reversing of the blood lymphopenia. The lack of MC-RANKL-controlled lymphocyte egress could either be compensated by other cells or overcome due to organ-based limits of cell hyperplasia. Importantly, although lymphocytes eventually reached the challenged ear skin at later time points, their delayed arrival was associated with a weaker inflammatory response. This suggests that the early phase of cell recruitment is critical for setting the inflammatory tone of the tissue. Even though total leukocyte numbers were increased at 48 h after DNFB challenge, the reduced infiltration at 24 h may have already limited the local production and amplification of inflammatory mediators. In particular, delayed T-cell entry could result in reduced availability of cytokines and chemokines that are needed to sustain inflammation and support the recruitment of additional effector cells. At the same time, mast cell-derived proteases may further shape this environment by processing or degrading cytokines and chemokines after degranulation. Since MC-specific RANKL deficiency did not affect mast cell degranulation, the phenotype is unlikely to reflect a general defect in granule release. Rather, our data suggest that MC-derived RANKL contributes to the timely recruitment of lymphocytes into inflamed skin and thereby influences the subsequent inflammatory cascade. In line with this, neutrophil infiltration into the challenged skin was also delayed, despite normal mobilization of neutrophils into the blood, indicating that the defect lies mainly at the level of tissue recruitment rather than systemic availability. Consequently, our data showed that early RANKL signaling induced by peripheral skin MCs is critical for timely lymphocyte egress from distant LN_in_, which in turn is crucial for proper onset of inflammation.

When investigating the long-range signaling mechanism, we observed an early rise in serum sphingosine-1-phosphate (S1P) levels just two hours after DNFB challenge, which was abolished in the absence of MC-derived RANKL. Importantly, injecting the same S1P amount i.v. at this time-point restored both the LN_in_ hyperplasia and blood lymphopenia, as well as lymphocyte infiltration into the skin to normal levels. It has been reported previously, that S1P is crucial for controlling lymphocyte egress from lymphoid tissues. S1P concentration is always higher in the blood and low in peripheral tissues. The S1P receptor 1 (S1P_1_) is internalized in circulating lymphocytes due to the high S1P concentration and re-expressed in S1P-poor tissues, allowing for a gradient promoting egress from secondary lymphoid organs to the blood (23). Adoptively transferred lymphocytes lacking the S1P_1_ can enter but not leave secondary lymphoid organs (33, 34). Deletion of the S1P lyase increases total S1P concentration in the LN and blocks lymphocyte egress. Similarly, deletion of both S1P kinases in lymphatic endothelial cells (LECs) blocks lymphocyte egress (35, 36). Therefore, the S1P spike in the blood after DNFB challenge in MC^wt^, but not MC^ΔRANKL^ mice, seems to be acting as a lymphocyte egress signal, while the early time-point indicates that preformed RANKL contained in the MC-granules may be responsible. Unexpectedly, intradermal injection of recombinant RANKL alone did not reproduce the rapid S1P burst observed 2 h after challenge. This indicates that RANKL is unlikely to act as an isolated trigger in vivo. Instead, the response may require prior tissue conditioning during sensitization or a synergistic effect with other mast cell-derived mediators. The local delivery of RANKL during focused degranulation can also be important. In this context, storage within mast cell secretory granules may stabilize RANKL, and enable its targeted and concentrated release into perivascular or lymphatic niches. Importantly, MC-RANKL could also act through direct MC interaction and communication with S1P controlling cells in the above niches. Thus, MC-derived RANKL appears to regulate early S1P release in a context-dependent manner rather than through simple soluble exposure alone. Importantly, we were able to show that *in vitro* cultured skin LECs respond to recombinant and MC-RANKL with the production of S1P. MC-RANKL could be acting on skin LECs or directly reaching the LN through the circulation. Thus, it is plausible that a MC-LEC communication axis is involved in controlling lymphocyte egress from distant LNs during skin inflammation.

Nevertheless, the contribution of additional cells cannot be excluded. Although inflammatory monocytes have been shown to raise LN S1P levels, thereby retaining lymphocytes in the tissue (37) and are recruited by MCs (12), the 2h time-point is too early for both a significant monocyte infiltration and their S1P synthesis. Therefore, we speculate the S1P spike to result from MC-RANKL effects on blood vessel endothelial cells (BECs). BECs express the receptor RANK, and RANKL stimulation promotes their survival and proliferation, thus inducing angiogenesis (38, 39). Moreover, BECs are one of the main controllers of serum S1P levels (40, 41). Given the ability of perivascular MCs to pass the vessel wall and degranulate directionally into the blood stream (15), RANKL contained in MC granules may activate BECs systemically even in distant blood vessels to immediately raise blood S1P concentration upon inflammatory insult. Moreover, S1P may be released by the MCs themselves, which are known as source of S1P, in an autocrine manner (42).

Conclusively, this study demonstrates that skin MC-derived RANKL is essential for lymphocyte egress from distant LN, via a mechanism involving S1P signaling. In absence of MC-derived RANKL, expanded lymphocytes do not egress from LNs in a timely manner upon skin inflammatory insult, resulting in massive LN hyperplasia and severe blood lymphopenia. Moreover, delayed lymphocyte egress and skin infiltration leads to attenuated CHS symptoms. Our results highlight the ability of peripheral MCs to act remotely and the importance of skin MC-derived RANKL in regulating lymphocyte egress and the initiation and progression of inflammation, with significant implications for the treatment of inflammatory diseases.

## Materials and methods

### Sex as a biological variable

Our study examined male and female mice, and similar findings are reported for both sexes.

### Mice

Mcpt5-Cre iDTR (Tg(Cma1-cre)ARoer x Gt(ROSA)26Sor^tm1(HBEGF)Awai^) and Mcpt5-Cre RANKL^FL/FL^ Tg(Cma1-cre)ARoer x B6.129-Tnfsf11^tm1.1Caob^/J mice were housed at the Central Animal Facility, Medical Faculty, OvGU Magdeburg, under specific pathogen-free conditions. All lines were on C57BL/6 background. In all experiments, mice were used at the age of 8–12 weeks. Littermates were used as controls. All procedures were in accordance with institutional guidelines on animal welfare and were approved by the Landesverwaltungsamt Sachsen-Anhalt, according to animal licenses 203.m-42502-2-1416_UniMD, and 203.m-42502-2-1732_UniMD. The Mcpt5-Cre line, Cre recombinase expression has been previously proven to be specific for connective tissue type MCs using a Cre-induced EYFP reporter line, while no expression was found in mucosal MCs as well as in other immune cells except a small fraction of NK cells(18).

### Contact hypersensitivity

Mice were sensitized with 100µl 0.5% DNFB (w/v) in acetone/olive oil (4:1) on the shaved back and challenged 6 days later with 20µl 0.2% DNFB (10µl each side of the ear). Vehicle controls were treated only with the solvent during the sensitization and challenge steps. Ear thickness was measured before and at several time points after challenge with an engineer’s micrometer (Mitutoyo, Takatsuku, Japan). Ear swelling as indicator of skin inflammation was calculated as percent increase compared to pre-challenge ear thickness.

For FITC-induced CHS, mice were sensitized by epicutaneous application of 100µl of fluorescein isothiocyanate (FITC; 0,5% w/v in dibutyl-phtalate/acetone 1:1) on the shaved back. 48h after sensitization, inguinal lymph node (LNs) were isolated and DCs were analyzed by flow cytometry.

### Adoptive transfer

CFSE labeled splenocytes were adoptively transferred in 37C pre-warmed, sterile PBS, by intravascular injection through the retro-orbital sinus.

### Cell preparation and flow cytometry

For preparation of skin cell suspensions, samples of ear skin were cut into small pieces and digested in RPMI 1640 (PAN Biotech, Aidenbach) containing, 0.025mg/ml Liberase TM, 396U/ml DNase I, and 0.5mg/ml hyaluronidase at 37°C, 1400 rpm for 1h. Samples were passed through a 40-mm sieve, and the cell suspension was washed twice with PBS.

For flow cytometry, isolated cells were incubated with an anti-FcγRIII/II antibody (clone 93) to block unspecific staining. Thereafter, cells were resuspended in PBS/2% BSA and 1–2 x10^6^ cells were stained with fluorochrome-conjugated monoclonal antibodies in 100µl staining solution for 15 min at 4°C. Cell suspensions were washed twice and were resuspended in 200µl PBS/2% BSA. Matched FMO controls were used to assess the level of background fluorescence in the respective detection channel. Flow cytometric analysis was performed on BD LSR Fortessa (BD, San Jose, CA) and analyzed with FlowJo Analysis Software (Ashland, OR).

### Preparation of single cell suspensions from spleen, mesenteric lymph nodes (mLN) and Peyers patches (PP) and analysis via flow cytometry

To prepare single cell suspensions, mLN were first cleared of adipose tissue, and mLN and PP were finely minced with a scalpel. Afterwards, Sp, mLN and PP were forced through 100μm filters and cells were centrifuged (400 x g, 5 minutes, RT). Splenocytes were subjected to ammonium-chloride-potassium lysis to remove erythrocytes, followed by quenching with RPMI 1640 (PAN Biotech) + 10% (v/v) FCS (Sigma). After centrifugation, final cell suspensions were filtered through a 40 μm filter. The following antibodies were used for staining: anti-mouse CD3 (17A2), CD8a (53-6.7), CD11c (N418), CD19 (6D5), CD45 (30-F11), CD62L (MEL-14), F4/80 (BM8), anti-mouse CD4 (GK1.5), CD11b (M1/70), CD44 (IM7), Ly6G (1A8), CD21/35 (eBio8D9), CD23 (B3B4), Foxp3 (FJK-16s), IgD (11-26c), and IgM (II/41). Cells were first stained for viability using Fixable Viability Dye eF780 for 20 min at 4°C. Afterwards, Fc receptors were blocked with anti-mouse CD16/32 (purified from 2.4G2 ATCC HB-197) for 10 min at 4°C prior to staining with surface antibodies for 30 min at 4°C. For intranuclear staining, samples were fixed and permeabilized using the Foxp3/Transcription Factor Staining Buffer Set for 30 min or overnight at 4°C, followed by a 60 min incubation with intranuclear antibodies at 4°C. Samples were measured on a LSRFortessa and analyzed with FlowJo 10. B cell subsets were gated according to Figure 2 in (43).

### Reverse transcriptase PCR (RT-PCR) and real-time quantitative PCR (RT-qPCR)

One inguinal lymph node was homogenized in 200 μL TRIzol using CK14 tubes and the Precellys 24 homogenizer according to the manufacturer’s protocol. Cultured PBMCs were resuspended in 1000 μL TRIzol. Following chloroform addition, total RNA was isolated according to the manufacturer’s protocol and quantified via NanoDrop spectrophotometry. RNA was subsequently reverse-transcribed using random hexamer primers and the Advantage RT-for-PCR kit according to the manufacturer’s instructions. RT-qPCR was performed using TaqMan Gene Expression Master Mix and TaqMan Gene Expression Assays: Ccl19 (FAM-MGB probe Mm00839967 g1), Ccl21 (FAM-MGB probe Mm03646971 gH), Cxcl9 (FAMMGB probe Mm000434946 m1), Cxcl10 (FAM-MGB probe Mm00445235 m1), Cxcl13 (FAM-MGB probe Mm00444534 m1), and Hprt (FAM-MGB probe Mm00446968 m1), Tnfsf11 (FAM-MGB probe Mm00441908 m1). All quantitative analyses were performed in triplicates on the qTOWER³ cycler and relative expression levels were determined using the ΔCT method.

### Whole mount LN staining and Microscopy

Mice were anaesthetized with isoflurane and injected i.v. with anti-CD31 AF594 mAb for the blood vessel stain. 5 min later, mice were sacrificed and inguinal LNs (LNs) were collected. The LNs were fixed overnight at 4C° in 1% PFA fixation buffer (0,7%w/v Na_2_HPO_4_, 1,5%w/v Lysine, 0,2%w/v NaIO_4,_ 1% PFA in ddH_2_O). Afterwards the LNs were incubated for 5 days in blocking buffer (0,02% goat serum, 1%BSA, 0,3% Triton X 100, 0,5µg/ml Streptavidin in PBS), followed by a 7 day primary antibody staining in blocking buffer, all in room temperature. T cell zones were stained with a rat monoclonal anti-mouse CD8 (53-6.7) Alexa Fluor 488 conjugate, B-cell zones with a rat monoclonal anti-mouse B220 (MAR-1) Alexa Fluor 647 conjugate and lymphatic structures with a rat monoclonal anti-mouse Lyve1 (ALY7) Brilliant Violet 421 conjugate. The antibody staining step was followed by a 1 day staining step with streptavidin BV421. Then, LNs were washed for one day in wash buffer (0,5% Thioglycerol, 0,2% Tritonx100 in PBS) at room temperature, followed by dehydration with 50% ethanol for 2h, 70% ethanol overnight and absolute ethanol for 2h. Subsequently the LNs were cleared in 100% ethyl cinnamate, by shaking at room temperature for 15 min, and stored in 100% until it was time for microscopy. Whole LNs were analysed by two-photon laser-scanning microscopy, fully submerged in ethyl cinnamate, as follows.

Two-photon laser scanning microscopy was performed with an upright, fixed stage Leica Stellaris 8 microscope with simultaneous detection via 4 external spectral non-descanned-detectors. Illumination was performed at 840 nm and 980nm with a MaiTai Deep See and an Insight X3 TiSa-laser. A Leica HC FLUOTAR L 16x/0, 60 IMM CORR VISIR objective was used to image the lymph nodes, which were immersed in an ethyl cinnamate filled glass petri dish. Bidirectional tile scans (25% overlap) of the lymph node were recorded as Z stacks (Δz = 5μm) encompassing the entire lymph node with an x-y resolution of 1024 x 1024 Pixel per single image at 600Hz Scanspeed with a line average of 4. Data were visualized as surface rendered z-stacks, or single z-planes. Raw data were reconstructed, analyzed and quantified using Imaris. For the blood vessel analysis, the Imaris filament tracer was used, while the T cell zones, B cell zones and lymphatic tissues were reconstructed with the surface tool. The volume of different LN compartments and blood vessel parameters, such as diameter and segment length, were calculated based on the surfaces and filaments.

For light-sheet microscopy, cervical LNs (cLNs) were isolated and fixed in 4% PFA in PBS for 2h at 4°C. All following steps were performed with permanent sample rotation: cLNs were washed in PBS and permeabilized in permeabilization buffer (20% DMSO, 0.3M glycine, 0.2% Triton-X, 2.5 µg/mL streptavidin in PBS) overnight at 4°C, followed by decolorization in 25% Quadrol (w/v) in dH2O overnight at room temperature. Afterwards cLNs were washed in PBS and blocked in blocking buffer (10% DMSO, 6% FCS, 0.2% Triton-X in PBS) overnight at 37°C. All staining steps were performed at 37°C. Primary antibody staining was performed in antibody buffer (5% DMSO, 3% FCS, 0.2% Triton-X, 0.05% NaN3 in PBS) with a rat anti-mouse monoclonal CD31 (MEC13.3) unconjugated antibody for 4 days. cLNs were washed 6x 30 min and overnight in washing buffer (10 µg/mL heparin, 0.2% Triton-X in PBS) and stained with an anti-rat DyLight 800 secondary antibody in antibody buffer for 4 days. The washing procedure was repeated and cLNs were stained with a rat anti-mouse monoclonal anti-mouse CD8 (53-6.7) Alexa Fluor 488 conjugate, rat anti-mouse monoclonal B220 (RA3-6B2) Alexa Fluor 647 conjugate, and Avidin TexasRed in antibody buffer for 3 days. After washing, cLNs were embedded in 1% agarose in dH2O followed by dehydration with 50% ethanol overnight, 70% ethanol for 8h and absolute ethanol overnight at room temperature. Finally, embedded cLNs were cleared in ethyl cinnamate (Eci) overnight at room temperature and stored in fresh Eci until imaging. Samples were imaged with a Miltenyi Biotec UltraMicroscope Blaze^TM^ light-sheet microscope. Images were acquired as tile scans and stitched with the Imaris stitcher (v 10.1.1). Image processing was performed with Imaris (v 10.1.1).

### Whole mount back skin staining and microscopy

For whole mount staining and light-sheet imaging of back skin samples, mice were sacrificed and a square of shaved skin was cut from the back. The skin samples were treated with hair removal cream for 1 min and washed in PBS. The fixation and permeabilization of the samples was performed as described above. Afterwards, the samples were further permeabilized in NaOH solution (0.1M NaOH + 0.01% Triton-X in PBS) for 20 min at 4°C. Samples were decolorized as described above, dehydrated by an increasing ethanol series, and bleached in bleaching solution (Methanol, DMSO, 30% H2O2 in a 1:4:4 ratio) overnight at 4°C. The samples were rehydrated and the whole mount staining was performed as described above. Primary antibody staining was performed with a rat anti-mouse monoclonal CD31 (MEC13.3) antibody and an Avidin Alexa Fluor488 conjugate for 7 days. Secondary staining was performed with an anti-rat DyLight800 secondary antibody and Sytox DeepRed nucleus dye for 6 days. Alexa Fluor 633 hydrazide was added to the staining solution for 24h. Finally, the samples were embedded and cleared as described above and imaged with a Miltenyi Biotec UltraMicroscope Blaze^TM^ light-sheet microscope.

### Dermal sheet staining and microscopy

To analyze MC numbers and localization in the ear skin, ears were isolated and split into dermal sheets. Samples were fixed overnight in 1% PFA in PBS at 4°C with mild shaking. Afterwards, samples were permeabilized on ice in a methanol/aceton solution (1:1) for 10 min per side, washed for 10 min in PBS, and incubated in 100 mM glycine for 10 min for fluorescence signal enrichment. After washing, blocking was performed with 0.5 µg/mL streptavidin in PBS for 60 min at RT. MCs were stained with an Avidin-TexasRed conjugate for 1h. For primary antibody staining, samples were washed 4x in PBS and blocked in blocking buffer (2% serum matching the secondary antibody, 50mM glycine, 0.05% Tween 20, 0.1% Triton X 100, 1% bovine serum albumin) for 60 min. Subsequently, samples were stained with a rat anti-mouse monoclonal (eBioV.7C7) endomucin-eFluor 660 conjugate, a mouse anti-mouse monoclonal (1A4) αSMA-eFluor660 conjugate, and a RANKL polyclonal antibody overnight at 4°C, followed by 4x washing in 0.1% Tween 20 in PBS. Secondary staining was performed with a goat anti-rat Alexa Fluor 647 conjugate and a goat anti-rabbit Alexa Fluor 488 conjugate in 0.1% Tween 20 in PBS for 1h at RT. After 3x washing, samples were stained with DAPI for 15 min at RT, washed 3x, and airdryed. Finally, the samples were mounted onto an object slide for microscopy in Vectashield Antifade Mounting Medium and the coverslip was sealed with nail polish. Imaging was performed within 1 to 4 days using an inverted confocal laser scanning microscope TCS SP8. For imaging of whole dermal sheets, illumination was performed via an HC PL APO 20x/0,75 IMM CORR CS2 oil dipping lens. The dermal sheets were imaged in tile scans and stitched afterwards with the Imaris stitcher (v 10.1.1). Images were recorded as 12-bit z stacks with a resolution of 512×512 pixels with depth of 200-250µm and a z spacing of 5 µm. For imaging of RANLK-positive MCs, samples were illuminated via the 40x oil objective. Images were recorded with a z spacing of 0.5 µm.

### Quantification of extravasation

To analyze lymphocyte extravasation from the blood vessels into the ear skin tissue (dermis) mice were challenged with 2% DNFB for 48h. Subsequently CD45 PE antibody was injected intravenously after mice were anaesthetized with an i.p. injection of ketamine 0,1 mg/kg, and Rompun 10 mg/kg. In this way, all lymphocytes located in the blood vessels were labeled with PE. 5 min after the injection of CD45 PE the mice were sacrificed, ears were collected, digested and a single cell suspension was prepared and stained for FACS. Lymphocytes that were located inside the blood vessels were therefore stained with the CD45 PE antibody and lymphocytes in the skin tissues were able to be separated by a second staining with CD45 PercP-Cy5.5 that was performed ex vivo.

### Culture of PCMCs

Peritoneal MCs were harvested by peritoneal lavage with 5ml ice-cold PBS and cultured in RPMI complete medium containing 10% FCS, 1% Pen/Strep, 50µM β-Mercaptoethanol, 1% sodium Pyruvate. The medium was supplemented with 10ng/ml IL-3 and 30ng/ml SCF. Medium was changed on days 2, 5 and 8 and the cells were used for reconstitution on day 10. From day 5 the SCF concentration was switched to 10ng/ml.

### PCMC degranulation assay

For quantification of MC degranulation, 1×10^5^ PCMCs were seeded in RPMI complete medium. For DNP/IgE stimulation, PCMCs were seeded in 150 ng/ml anti-DNP IgE for sensitization. On the next day, sensitized PCMCs were stimulated with 100ng/ml DNP-BSA and untreated PCMCs with 500 ng/ml calcium ionophore A23187, 500 µM ATP or 50 ng/ml IL-33 in a total volume of 125 µl RPMI complete medium for 10 min, respectively. Then, cell culture plate was placed on ice for 5 min and centrifuged (365 g, 5 min, 4°C), before cells were stained for flow cytometry as described above.

### Mast cell depletion and local reconstitution of ear skin

In Mcpt5-Cre iDTR mice, MCs were locally depleted in the ear skin by a single injection of 100ng of DT in 20µl sterile PBS. 5 days after MC depletion the ears were reconstituted with 1 million PCMCs in 20µl of sterile PBS per ear. The cells were allowed to repopulate the tissue for 2 weeks before the induction of CHS.

### Automated multidimensional fluorescence microscopy by multi-epitope-ligand cartography (MELC)

MELC was performed as described previously (1). Briefly, peripheral lymph nodes from RANKL WT and KO mice were embedded into in Tissue-Tek® O.C.T.™ Compound, frozen by liquid nitrogen and stored at -80°C. 10 µm cryo-sections adhered to silan-coated cover slides were fixed with 2 % paraformaldehyde and permeabilized with 0.2 % Triton-X-100 and blocked with 1 % BSA+ 30% NGS. The tissue samples were transferred to an inverted wide-field fluorescence microscope (Leica DMi8, 20x air lens NA 0.80). The automated cyclic robotic process started with the incubation of the first fluorescently labelled antibody (tag). After a series of washing steps, the fluorescence signals and a corresponding phase contrast image were acquired by a scientific CMOS camera (Orca Fusion, Hamamatsu). The specific signal of the given tag was removed by bleaching of the fluorescent dye followed by recording of post-bleaching fluorescence signals and repetition of incubation-imaging-bleaching-cycle. The appropriate working dilutions, incubation times, and positions within the MELC experiment were validated systematically using conditions suitable to MELC. The series of fluorescence images produced by each tag were aligned with a accuracy of 0.1 pixels using the corresponding phase contrast images. Illumination faults of the images were corrected using flat-field correction. Post-bleaching images were subtracted from the following fluorescence tag images. Section artifacts were excluded as invalid by a manual mask-setting process. For cell segmentation we used a summed image of the marker CD45 and CD54 and recall the general model of Cellpose (44) within our MATLAB analysis workflow. The CellProfiler (45)(software was used to generate a mask for the FRC region, defined as Gp38+ and CD31- cell using the signals of Gp38 and CD31. For each cell the mean fluorescent intensity and the smallest distance to the reference region FRC was calculated. The resulting matrix of intensities and distances were exported into an FCS file and uploaded to the online cytometry analysis platform “cytobank.org” for multi-parametric analysis.

### Visualizing RANKL in MCs *ex vivo* and *in vitro*

Mouse ear skin from Mcpt5-Cre-iDTR mice (Cre⁺ and Cre⁻) was harvested, embedded, frozen, and sectioned. Sections were fixed in 1% PFA/PBS overnight at 4°C in a humidified chamber, followed by permeabilization in methanol/acetone (1:1) for 2 min on ice. After washing with PBS, MCs were streptavidin blocked (0.5 µg/mL, 60 min) followed by staining with labeled Avidin–Alexa Fluor 488 (50 ng/mL, 60 min, room temperature). Sections were then blocked for 60 min at room temperature and incubated with anti-RANKL PE primary antibody (1:80) overnight at 4°C. After washing (PBS, 0.1% Tween-20), sections were incubated with goat anti-rat Alexa Fluor 647 secondary antibody (1:500) for 1 h at room temperature. Slides were washed, air-dried, and mounted for imaging.

For in vitro experiments, MCs (Cre+ and Cre- from Mcpt5-Cre-iDTR mice) were isolated, cultured, and cytospun onto glass slides (1×10⁵ cells per slide). Cells were stained using the same protocol as described for skin sections. Images were obtained with an 63x objective and z-stacks spacing of 0.3 µm.

### Dermal skin cytokine and serum S1P quantification

To enable the detection of cytokines, mice were sensitized and challenged. Then the ear collection at 48h and serum collection at 2, 4, 6, 12, 24 and 48 hours too place. Ear skin samples were shock frozen in liquid nitrogen, suspended in serum-free medium and mechanically homogenized using a tissue homogenizer mill with stainless steel or ceramic beads. Cytokines of the ear skin were quantified, using a Legendplex assay (Biolegend) according to manufacturer’s instructions, and were normalized to skin biopsy weight. S1P was quantified using a reverse inhibition ELISA (Biomatik) according to manufacturer’s instructions.

### Isolation and culturing of

Lymphatic vessel endothelial cells (LECS). LECs were isolated from C57B/6J mice. The mice were sacrificed for scientific purposes using CO2 inhalation, according to the IMKI-TWZ-ADU-1-25. The ears of each mouse were collected, separated into dermal sheets and minced into 1 mm2 pieces, and enzymatically digested with 198U/ml DNAse, 0.03125 mg/ml Liberase and 0.25 mg/ml Hyaludonidase in un-supplemented RPMI, vigorously shaken at 37o C for 1 hour. The cell suspension was filtered through a 100 μm filter, and the cells were pelleted at 300g for 5 minutes at 4oC. The supernatant was discarded, and the cells were resuspended in MACs buffer (PBS, 2mM EDTA, 0.5% w/v BSA) containing 10 μl/ml LYVE-1-biotin, 15 μl/ml CD45-eFlour450 and 15 μl/ml CD31-AF488 antibodies, and incubated for 15 minutes at 4oC. The primary antibodies were washed with MACs buffer, and the cells were pelleted at 300g for 5 minutes at 4oC, resuspended in MACs buffer containing 10 μl/ml Streptaviding-AF647, and incubated for 30 minutes at 4oC. Following the incubation, the cells were washed with MACs buffer, pelleted, resuspended in fresh MACs buffer, and sorted using ARIA III Cell Sorter (BD). From the CD45 negative population, the population that was double positive for LYVE-1 and CD31 was collected. After the sorting the cells were pelleted and resuspended in high glucose DMEM (PAN Biotech) supplemented with 20% FCS,100U/ml penicillin, 100 μg/ml streptomycin, 2mM L-glutamine, 20 mM HEPEs, 1% minimally essential amino acids, 1mM sodium pyruvate, 50 μM 2-mercaptoethanole, 75 μg/ml endothelial growth factor, 1 U/ml of heparin, 10 μg/ml amphotericin B and 10 μg/ml Fungin. The cells were seeded in well on a 24 well plate coated with 2% gelatin and incubated at 37o C in a 5% CO2 environment. The media was replaced the day after the sorting, and afterwards every 2 to 3 days. Amphotericin B was removed from the culture after 3 to 4 days, while Fungin was after a week.

After the cells sorting, the cells were allowed to grow to 80% confluence, before they were sculptured to a larger container. When subcultured, the cells were washed with PBS to remove protein residues and incubated with 0.5% trypsin EDTA at 37o C in a 5% CO2 environment until they detached from the plate. The trypsin was neutralized with complete EC medium, and the cells were pelleted at 300g for 5 minutes at 4oC. To remove the trypsin and gelatin residues, the supernatant was discarded and the cells were resuspended in complete EC medium, before transferred to a fresh container coated with 2% gelatin. The cells were subcutured 6 times before being characterized or used for experiments.

#### Software

For visualization, 2-photon microscopy raw data were processed and reconstructed using Imaris V10.1 (Bitplane/Zurich/Switzerland). Analysis of flow cytometry raw data was performed using FlowJo Analysis Software V10 (FlowJo LLC, Ashland, OR). Graphic data analysis was done with the OriginPro software (OriginLab Corporation, Northampton, MA) and GraphPad Prism 10 software, (Graphpad software, San Diego, CA). Statistical analysis was performed with GraphPad Prism 10.

#### Statistics

Statistical analysis was performed with the Mann-Whitney U test, when comparing two experimental groups, or otherwise using ANOVA. The respective non parametric tests were used in the case where a non-Gaussian distribution was found. All graphs are depicted as mean and SD. **** p < 0.0001; *** p < 0.001; ** p < 0.01; * p < 0.05.

#### Study approval

All procedures were in accordance with institutional guidelines on animal welfare and were approved by the Landesverwaltungsamt Sachsen-Anhalt (42502-2-1416UniMD).

## Data availability

All data are available in the Supporting Data Values file and are available upon request.

## Acknowledgements

We sincerely thank all members of the Dudeck laboratory for discussions and support. Special thanks go to expert technical support in method establishment by M. Voss, H. Edler, J. Fröbel, Camilla Merten, Jeannette Seger, Nicole Jüling and O. Biskou. Moreover, we thank A. Müller and R. Hartig from the Multiparametric Bioimaging and Cytometry (MPBIC) platform for support with microscopy and cytometry approaches. This work was funded by the Deutsche Forschungsgemeinschaft (DFG, German Research Foundation: Project-ID 361210922/RTG2408/TP4, DU1172/8-1, and DU1172/9-1 to AD, and supported by the Health Campus for Inflammation, Immunology and Infectiology of the Medical Campus of the Otto-von-Guericke University (GC-I3). Disclosure of potential conflict of interest: L. Philipsen. is affiliated with BioDecipher GmbH, a company which develops systems for multiplexed fluorescence microscopy (MELC).

## Author Contributions

KKD and AD conceived and designed the study, analyzed and discussed data and wrote the manuscript. KKD, WU, AEB, AH, VD, LK, TS, CH, LS, NJN and JD performed experiments and analyzed data. AJM, SK, TS and SF provided significant guidance and expertise during the revision process, including advice on experimental design and critical review of the manuscript’s content. All authors revised the manuscript and approved the final version of the manuscript.

