## Supplementary material for "Mast cells initiate lymphocyte egress from distant lymph nodes upon skin inflammation via a RANKL–sphingosine-1-phosphate axis": Suplemental Figures and Tables

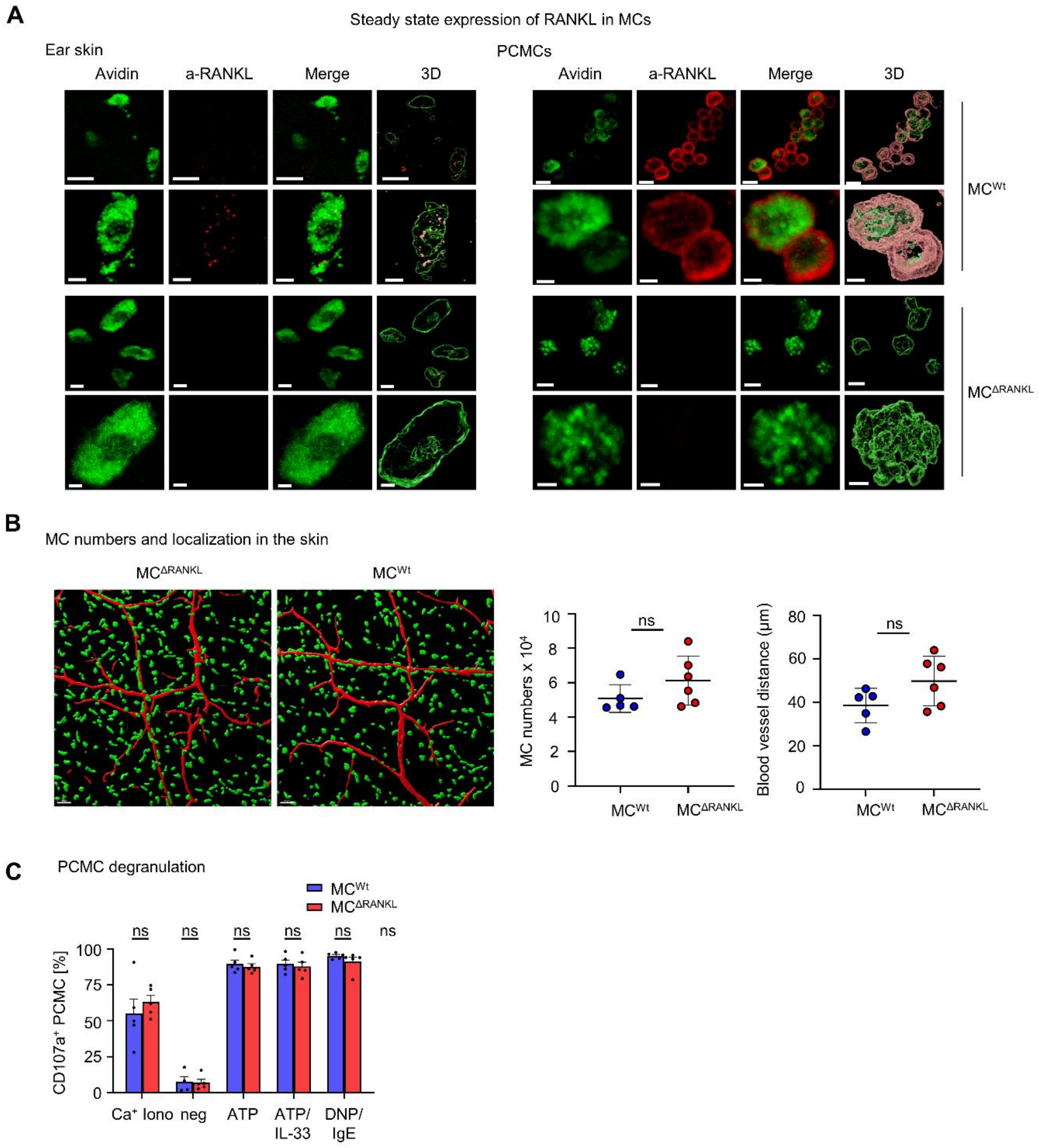


**Supplemental Figure 1: Absence of MC-derived RANKL does not affect MC degranulation and skin localization.** (**A**) RANKL expression in $MC^{\Delta RANKL}$ and $MC^{Wt}$ mice in steady state ear skin sections (left panel) and peritoneal cultured mast cells (PCMCs) (Right panel) (**B**) Whole mount ear dermal sheet microscopy of $MC^{\Delta RANKL}$ and $MC^{Wt}$ mice. MC numbers and distance to the blood vessels were quantified with Imaris software. (**C**) PCMCs were cultured and degranulation was induced with different stimuli. All graphs are depicted as mean and SD. **** $p < 0.0001$; *** $p < 0.001$; ** $p < 0.01$; * $p < 0.05$.

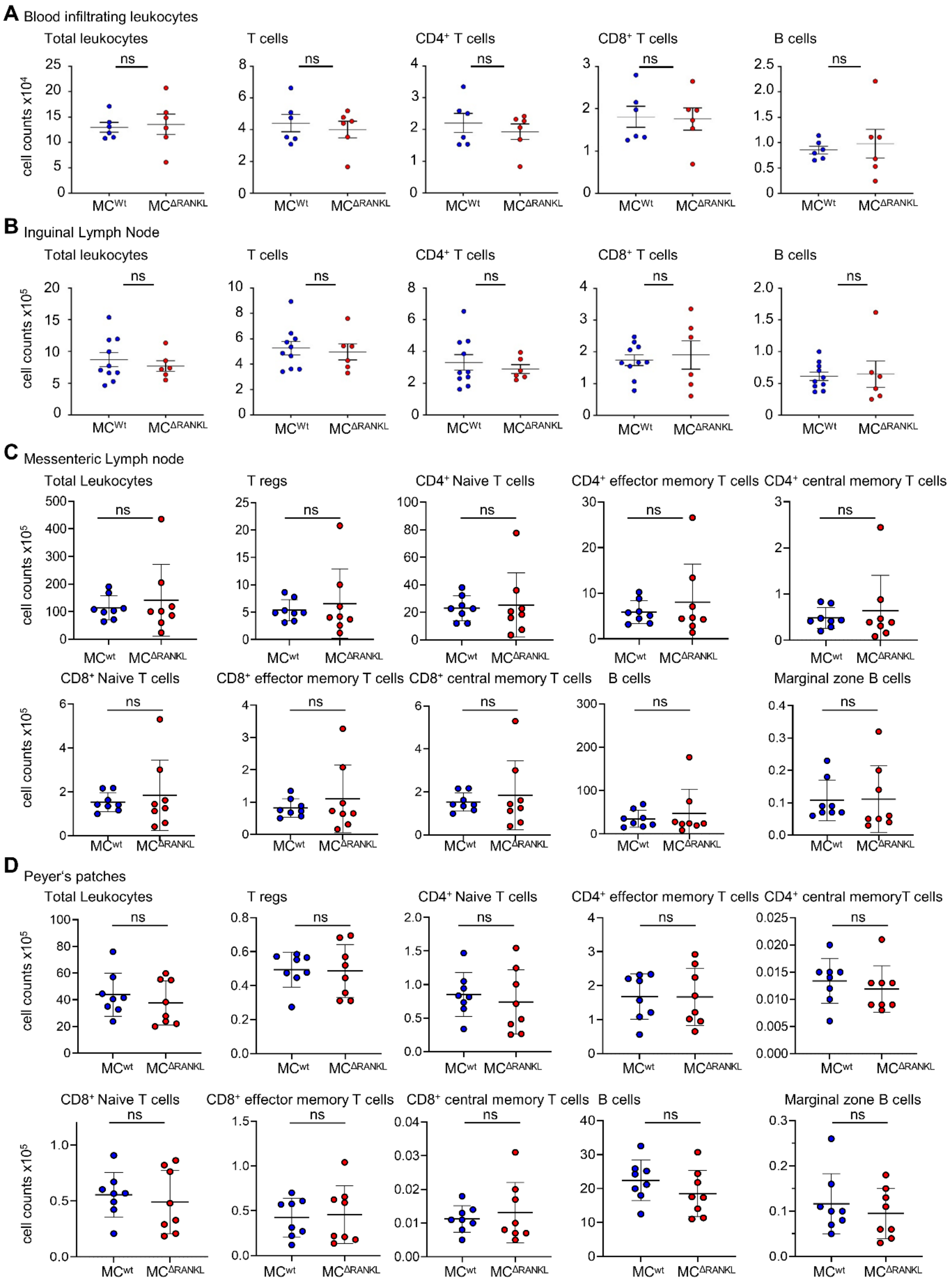


**Supplemental Figure 2: Absence of MC-derived RANKL does not affect steady state leukocyte populations.** Blood and LN lymphocyte numbers under physiologic conditions were analysed by flow cytometry. (**A**) Blood lymphocyte numbers per 100µl of blood. (**B**) Total numbers of lymphocytes in $LN_{in}$. (**C**) Total numbers of lymphocytes in mesenteric LNs. (**D**)Total lymphocyte numbers in Peyer's Patches. All graphs are depicted as mean and SD. **** $p < 0.0001$; *** $p < 0.001$; ** $p < 0.01$; * $p < 0.05$.

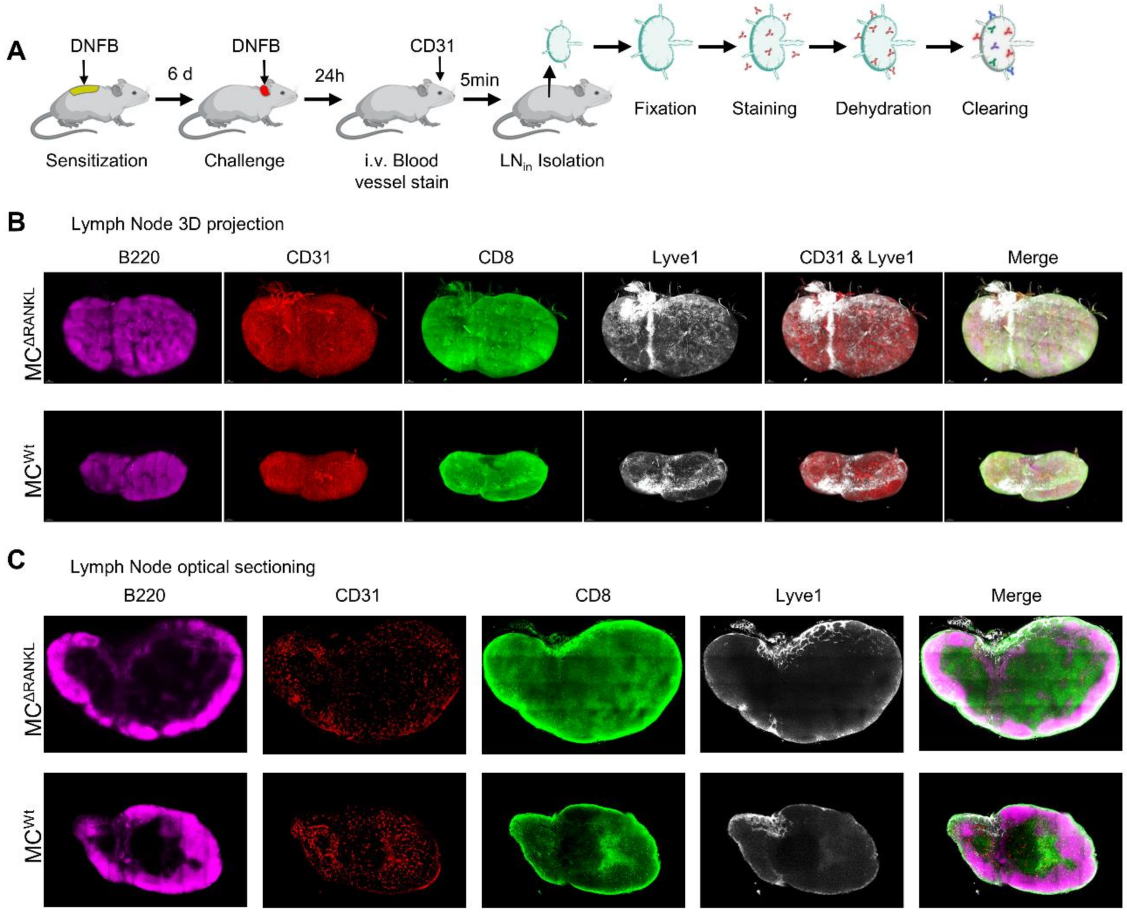


**Supplemental Figure 3: $LN_{in}$ hyperplasia in $MC^{\Delta RANKL}$ mice was investigated by advanced whole mount 3D microscopy.** (**A**) Mice were sensitized on the back skin and after 6 days challenged on the ear skin. 24h after challenge $LN_{in}$ were collected, stained with antibodies and cleared. LNs were stained with B220 for the B cell compartments, i.v. CD31 for the blood vessels, CD8 for the T cell compartments and Lyve1 for the lymphatic vessels and visualized with intravital two photon microscopy. (**B**) 3D lymph node projection. (**C**) Optical sections of $LN_{in}$ were taken from the midpoint of the imaging plane.

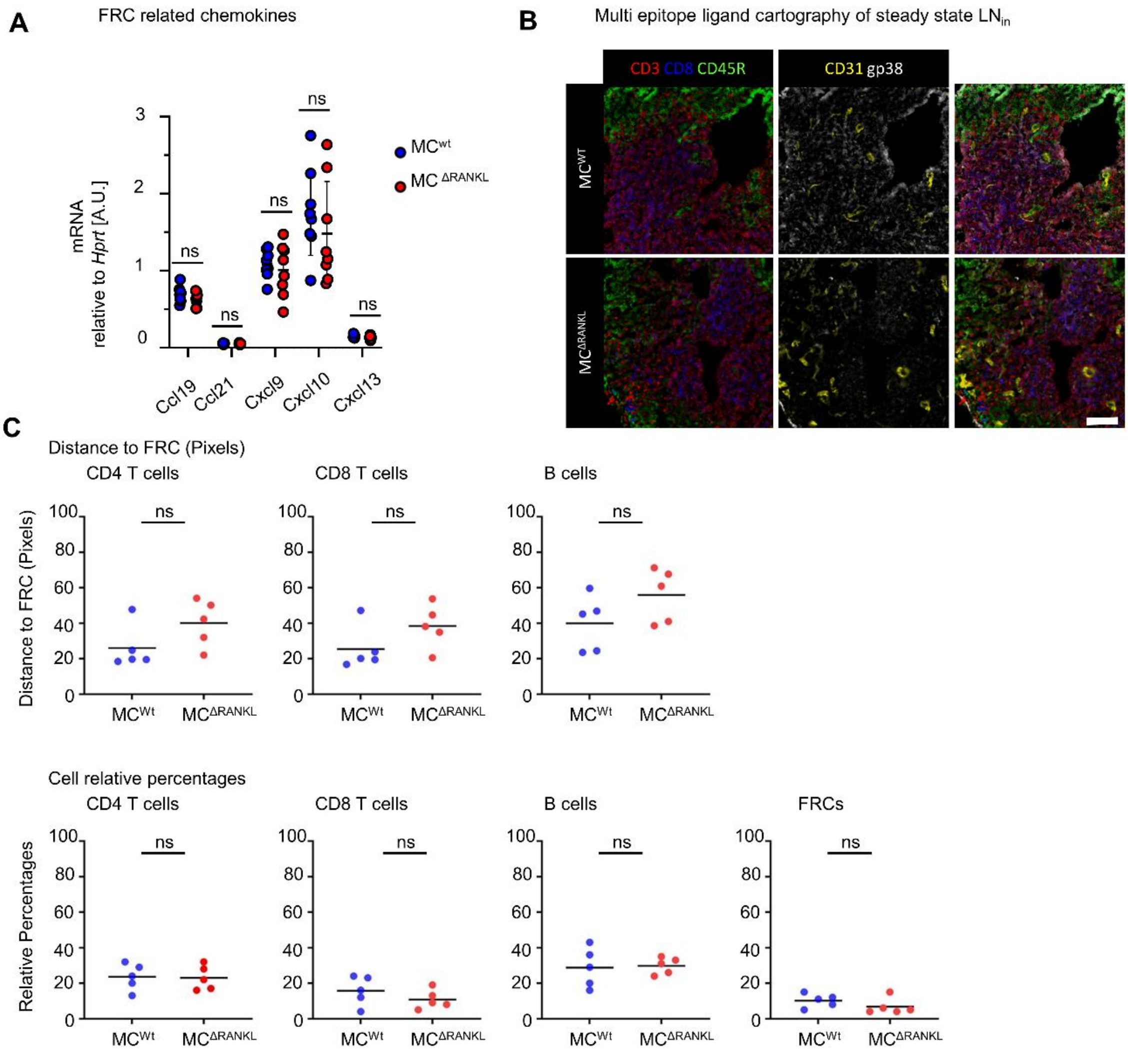


**Supplemental Figure 4: Absence of MC-derived RANKL does not affect LN fibroblastic reticular cell localization and function in the steady state.** (**A**) FRC relevant chemokine mRNA expression from whole inguinal LNs of $MC^{\Delta RANKL}$ and $MC^{Wt}$ mice. (**B**) Multi epitope ligand cartography was performed to identify FRCs, CD4 T cells, CD8 T cells and B cells in inguinal LN sections and (**C**) their distances from FRCs were calculated.

| REAGENT or RESOURCE | SOURCE | IDENTIFIER |
|---|---|---|
| Antibodies | | |
| Hamster monoclonal anti-mouse CD3 (145-2C11) PE-Cy7 | Biolegend | Cat# 100320; RRID AB_312685 |
| Hamster monoclonal anti-mouse CD3 (145-2C11) PE | eBiosciences | Cat# 12-0033-82; RRID AB_842786 |
| Rat monoclonal anti-mouse CD4 (GK1.5) Percp-efluor-710 | eBiosciences | Cat# 46-0041-82; RRID AB_11150050 |
| Rat monoclonal anti-mouse CD4 (GK1.5) AF488 | eBiosciences | Cat# 53-0041-82; RRID AB_469892 |
| Rat monoclonal anti-mouse CD8 (53-6.7) BV510 | Biolegend | Cat# 100752; RRID AB_2563057 |
| Rat monoclonal anti-mouse CD8 (53-6.7) AF488 | Biolegend | Cat# 100723; RRID AB_389304 |
| Rat monoclonal anti-mouse CD25 (PC61) BV421 | Biolegend | Cat# 102043; RRID AB_2562611 |
| Rat monoclonal anti-mouse CD44 (IM7) PE | Biolegend | Cat# 103008; RRID AB_312959 |
| Rat monoclonal anti-mouse B220 (MAR-1) AF647 | BD Pharmingen | Cat# 557683; RRID AB_396793 |
| Rat monoclonal anti-mouse B220 (MAR-1) FITC | Biolegend | Cat# 103206; RRID AB_312991 |
| Rat monoclonal anti-mouse Ly6G (1A8) PE-Cy7 | Biolegend | Cat# 127 618; RRID AB_1877261 |
| Rat monoclonal anti-mouse Ly6G (1A8) APC | Biolegend | Cat# 127614; RRID AB_2227348 |
| Rat monoclonal anti-mouse F4/80 (BM8) APC-efluor780 | eBiosciences | Cat# 47-4801-82; RRID AB_2735036 |
| Rat monoclonal anti-mouse CD11b (M1/70) Efluor-450 | eBiosciences | Cat# 48-0112-82; RRID AB_1582237 |
| Rat monoclonal anti-mouse Lyve1 (ALY7) Biotin | eBiosciences | Cat# 13-0443-82; RRID AB_1582237 |
| Rat monoclonal anti-mouse CD31 (390) | Biolegend | Cat# 102432; RRID AB_2617017 |
| Rat monoclonal anti-mouse CD31 (MEC13.3) | BD | Cat# 550274 RRID AB_393571 |
| Rat monoclonal anti-mouse CD45 PE | Biolegend | Cat# 103106; RRID AB_312971 |
| Rat monoclonal anti-mouse CD45 efluor-450 | eBiosciences | Cat# 48-0451-82 RRID AB_312971 |
| Rat monoclonal anti-mouse CD45 PercP-Cy5.5 | Biolegend | Cat# 103132; RRID AB_893340 |
| Goat polyclonal anti-rabbit AF488 | ThermoFisher | Cat# A-11006 |
| Goat polyclonal anti-rat AF647 | Jackson ImmunoResearch | Cat# 112-605-003 |
| Rabbit anti-mouse polyclonal RANKL | antibodies online | Cat# ABIN668556 |
| Mouse anti-mouse monoclonal αSMA (1A4) eFluor660 | eBioscience | Cat# 50-9760-82 RRID AB_2574362 |
| Rat monoclonal anti-mouse endomucin (eBioV.7C7) eFluor660 | eBioscience | Cat# 50-5851-80 RRID AB_11220069 |
| Anti-rat polyclonal DyLight 800 | invitrogen | Cat# SA5-10032 RRID AB_2556612 |

| | | |
|---|---|---|
| Mouse monoclonal anti-mouse Dinitrophenyl (DNP) IgE | Sigma-Aldrich | Cat# D8406; AB_259249 |
| Hamster monoclonal anti-mouse FcεRI (MAR-1) AF647 | Biolegend | Cat# 134309; AB_1626097 |
| Rat monoclonal anti-mouse CD117 (ACK2) BV421 | Biolegend | Cat# 135124; AB_2562237 |
| Rat monoclonal anti-mouse CD107a (1D4B) PE | eBiosciences | Cat# 12-1071-82; AB_657554 |
| Bacterial and Virus Strains | | |
| n/a | | |
| Biological Samples | | |
| n/a | | |
| Chemicals, Peptides, and Recombinant Proteins | | |
| DNFB (1-fluoro-2,4-dinitrobenzene) | Sigma-Aldrich | Cat# D1529-25ML |
| Hyaluronidase | Sigma-Aldrich | Cat# H3506 |
| Liberase™ | Roche | Cat# 5401119001 |
| DNase I | Roche | Cat# 10104159001 |
| Olive oil | Sigma-Aldrich | Cat# O1514-100ML |
| Aceton | Fisher Scientific/J.T.Baker | Cat# 8002 |
| Streptavidin BV421 | Biolegend | Cat# 405226 |
| Sphingosine 1 Phosphate | Sigma Aldrich | Cat# 73914-1MG |
| Thioglycerol | Sigma-Aldrich | Cat# 88640-100ML |
| Ethyl Cinnamate | Sigma-Aldrich | Cat# 199179-92-5 |
| CellTrace™ CFSE Cell Proliferation Kit | Thermo Fisher | Cat# C34554 |
| Sytox DeepRed nucleus dye | invitrogen | Cat# S34859 |
| AF633 hydrazide | life technologies | Cat #A30634 |
| Streptavidin | Jackson ImmunoResearch | Cat# 016-000-113 |
| N,N,N′,N′-Tetrakis(2-hydroxypropyl)ethylendiamin (Quadrol) | Tokyo Chemical Industry | Cat #T0781 |
| DAPI | Sigma-Aldrich | Cat# 32670-5MG-F |
| Avidin TexasRed | MolecularProbes | Cat# A820 |
| Avidin AF488 | MolecularProbes | Cat# A21370 |
| Hair removal cream | Veet | Cat# 3018119 |
| Vectashield antifade mounting medium | Biozol | Cat# VEC-H-1000 |
| 2,4-Dinitrophenylated (DNP)-BSA (Albumin from Bovine Serum) | Thermo Fisher | Cat# A23018 |
| Calcium Ionophore A23187 | Sigma Aldrich | Cat# C7522 |
| Adenosine 5′-triphosphate disodium salt trihydrate (ATP) | Roche | Cat# 10519979001 |
| Mouse IL-33 Recombinant Protein | PeproTech | Cat# 210-33 |
| Zombie NIR™ Fixable Viability Kit | Biolegend | Cat# 423106 |
| Critical Commercial Assays | | |
| Legendplex mix and match subpanel | Biolegend | Cat# CLT100520 |
| Sphingosine-1-Phosphate (S1P) ELISA Kit | Biomatik | Cat# EKU11440 |
| Deposited Data | | |
| n/a | | |

| | | |
|---|---|---|
| Experimental Models: Cell Lines | | |
| n/a | | |
| Experimental Models: Organisms/Strains | | |
| Mcpt5-Cre x iDTR (*Mus musculus*) (Tg(Cma1-cre)ARoer x Gt(ROSA)26Sor$^{tm1(HBEGF)Awai}$) | Central Animal Facility, OvGU Magdeburg | |
| Mcpt5-Cre x RANKL$^{Fl/Fl}$ (*Mus musculus*) Tg(Cma1-cre)ARoer x B6.129-Tnfsf11$^{tm1.1Caob}$/J | Central Animal Facility, OvGU Magdeburg | |
| Software and Algorithms | | |
| Imaris (for Windows) Version 9.5 | Oxford Instruments, Zurich, Switzerland | N/A |
| FlowJo$^{TM}$ Software (for Windows) Version 10 | FlowJo Inc. | N/A |
| GraphPad Prism (for Windows) Version 9.02 | Graphpad Software | N/A |
| ORIGIN-Pro | OriginLab Corporation | N/A |

Supplemental table 1: Catalogue of all reagents, mouse models and software used in this paper.
